# On the diversity of nematode antagonists in an agricultural soil, and their steerability by root-knot nematode density and cover crops

**DOI:** 10.1101/2024.09.27.615391

**Authors:** Sara G. Cazzaniga, Philippe Belliard, Joris van Steenbrugge, Sven van den Elsen, Carin Lombaers, Johnny Visser, Leendert Molendijk, Jose G. Macia-Vicente, Joeke Postma, Liesje Mommer, Johannes Helder

## Abstract

Plant-parasitic nematodes are harmful pathogens for many agricultural crops. Within this category, root-knot nematodes (RKN, *Meloidogyne* spp.) are worldwide regarded as the most impactful because of their wide geographical distribution and their polyphagous nature. Host plant resistances against RKN have been successfully introduced in a few crops only. As the use of nematicides is becoming increasingly restricted because of environmental and human health concerns, there is a need for alternative strategies to control RKN. One such approach is the stimulation of local nematode antagonists. We investigated this in an experimental field setting with two main variables: density of the Columbia root-knot nematode *Meloidogyne chitwoodi*, and the type of cover of crop. For each of the three *M. chitwoodi* densities, the effects of ten cover crop treatments were tested on both the resident (DNA) and the active (RNA) fractions of the bacterial and fungal communities. In our analyses, we focussed on changes in the abundance of plant-parasitic nematode antagonists. From the eight bacterial and 26 fungal genera known from global literature to harbour potential antagonists of plant-parasitic nematodes, we detected respectively five and 14 genera in our agricultural field. Among the bacterial genera, four included bacterial species for which nematode antagonism has been documented. The fungal genera included facultative nematode parasites (*e.g*., *Arthrobotrys* spp.), endophytes strengthening host defences (e.g., *Acremonium* spp.), as well as multiple obligatory nematophagous species. This study revealed that conventionally managed arable fields may harbour an unexpectedly high diversity of nematode antagonists. Multiple antagonists were stimulated by cover crops in a cover crop-specific manner, and, to a lesser extent, by increased RKN densities. The richness in putative nematode antagonists did not translate into *M. chitwoodi* suppression probably because most antagonists have a facultative nematophagous lifestyle and will only predate nematodes under poorer nutritional conditions.

## 1. Introduction

Global crop yield is severely limited by pests and diseases, causing up to 40% losses on a wide range of economically important crops worldwide (Savary et al., 2019). Plant-parasitic nematodes alone account for an estimated loss of US$ 173 billion every year (Elling, 2013), which makes the implementation of disease control measures a necessity in agriculture. Among the plant-parasitic nematodes, root-knot nematodes (RKN, *Meloidogyne* spp.) are most impactful worldwide (Jones et al., 2013) affecting almost every major crop.

Broadly speaking, three types of measures have been developed to control plant-parasitic nematodes. First, crop rotation is the practice by which the growing of susceptible plant species is alternated with non-host plants over time. Discontinuous presence of hosts has been shown to effectively reduce densities of several plant-parasitic nematode species (Azlay et al., 2023). However, crop rotation is barely effective against RKN as all agronomically relevant RKN species are highly polyphagous. Second, breeding for host plant resistances has been very effective in case of several major plant-parasitic nematode species. For over 60 years the dominant resistance *Mi1.2* gene has been effective in reducing the damage caused by the three ‘tropical’ root-knot nematode species (*M. incognita, M. javanica* and *M. arenaria*) in tomato (Milligan et al., 1998). However, the transferability of this R gene was shown to be limited to close relatives of tomato (Goggin et al., 2006). The scarceness of effective R genes and their limited transferability to other crops has limited the broadening of the application of this type of control. Third, a range of synthetic nematicides has been introduced to control plant-parasitic nematodes. General soil fumigants and inhibitors of the neurotransmitter acetylcholinesterase have been widely applied to control plant-parasitic nematodes for decades. Mainly due to their strong negative side effects on non-target organisms as well as their serious risks for human health (Heckel, 2012; Gill and Garg, 2014; Prashar and Shah, 2016), many of these nematicides have been phased out lately. Hence, due to the limited applicability of crop rotation, the limited availability of effective host plant resistance genes, and the widely supported endeavour to reduce the application of synthetic nematicides, there is a need for the development of alternative sustainable nematode control practices.

In soil, a diversity of antagonists against plant-parasitic nematodes has been identified (for reviews see, e.g., Li et al., 2015 and Topalovic et al., 2020). For example, the bacterial genus *Pasteuria* has diversified in a number of species that all parasitize on (mainly) plant-parasitic nematodes, with species-specific food preferences (Ciancio, 2018). Among the fungal nematode antagonists, *Arthrobotrys oligospora* (= *Orbilia oligospora*) can produce adhesive trapping nets in the presence of nematodes (Nordbring-Hertz and Mattiasson, 1979) and the closely related species *A. dactyloides* forms three-celled constricting rings to catch plant-parasitic nematodes (Higgins and Pramer, 1967). Yet another mechanism is employed by the fungus *Pochonia chlamydosporia* which parasitizes nematode eggs and additionally triggers defence responses in the plant root (Gouveia et al., 2023). Numerous attempts to introduce mass-produced nematode antagonists in soil have not resulted in successful agricultural applications. Generally speaking, the chances of successfully introducing microbial antagonists into a highly competitive environment such as soil are low (Giuma and Cooke, 1974; Jaffee et al., 1996; Stirling, 2011). As an alternative, it has been proposed to identify factors in cropping systems that could be used to support local nematode antagonists. Apart from organic amendments and mulches, cover crops were proposed as a means to support the growth and the activity of nematode antagonists (Stirling, 2011)

Cover crops are non-economic crops grown in between main crops to minimise nutrient leaching and soil erosion, and to increase the soil organic matter content (Blanco-Canqui and Ruis, 2020). A substantial number of plant species can be grown as a cover crop provided that they grow well outside the main growing season and as long as they can be terminated easily. Recent evidence has shown that cover crop species simulate or repress specific fractions of the soil microbial communities (Cazzaniga et al., 2023a). Such manipulation of the soil microbial community could be instrumental in the defence against plant pathogens (Berendsen et al., 2012; Philippot et al., 2013). We hypothesize that cover crops can be applied to boost the nematode antagonistic potential of soils by increasing the abundance and the activity of microbes antagonistic to plant-parasitic nematodes.

In this study, we performed an experiment in a field with known presence of *M. chitwoodi*. We focused on the Columbia root-knot nematode *Meloidogyne chitwoodi*, a highly polyphagous obligatory plant parasite that poses a significant threat to crops in temperate regions in both the northern and southern hemisphere (O’Bannon et al., 1982; Azlay et al., 2023). Because of its broad host range, *M. chitwoodi* cannot be effectively controlled by crop rotation, and although host plant resistances have been identified (Mojtahedi et al., 1995), they have not been introduced yet in commercial varieties of any main crop (Teklu et al., 2023). Hence, we investigated whether it would be possible to stimulate local nematode antagonists to control *M. chitwoodi* in an experimental field setting using cover crops.

First, we experimentally modified *M. chitwoodi* densities by growing four different grass species with different host status for this pathogen. Subsequently, we grew a range of cover crop monocultures and cover crop mixtures in plots with distinct nematode densities. With this setup we aimed to assess the impact of two main variables, cover crop identity and *M. chitwoodi* density, on the resident (DNA-based) and active (RNA-based) fractions of the bacterial and fungal communities. We focused on changes in genera known to comprise putative nematode antagonists.

The overall aim of this study is to reveal whether the growth of specific plant species or distinct levels of *M. chitwoodi* can strengthen the local nematode-suppressive potential of soils. To this end, we addressed the following questions: 1) Do cover crops affect the abundance and/or activity of local plant-parasitic nematode antagonists? 2) Does exposure to distinct *M. chitwoodi* levels affect abundance and/or activity of these antagonists? 3) Do cover crops and/or *M. chitwoodi* levels affect local plant-parasitic nematode antagonists generically or in a cover crop- and/or *M. chitwoodi* level-specific fashion?

## 2. Materials and methods

### 2.1 Experimental field set-up

The experimental field was located in Vredepeel (Limburg, the Netherlands), an experimental field station of the Field Crops unit (WUR-FC) of Wageningen University and Research. This field was characterized by sandy soil (1% clay, 8% silt and 87% sand) with an organic matter content of ≈ 4% (4.1 - 4.4%), and a pH of around 6 (5.4 - 6.1). Our experiment was embedded in a larger experiment by WUR-FC aimed at assessing the host plant status of a selection of arable crops and cover crops in an arable field naturally infested with *M. chitwoodi* (Visser et al., 2022). To generate four different initial population densities of *M. chitwoodi*, four pre-crops belonging to the Poaceae family with distinct host status for this plant-parasitic nematode were selected (Table 1). Pre-crops were sown in August in essentially three blocks each subdivided into four rectangular strips (6 x 42 m) (Figure S1). Two blocks (blocks 2 and 3) were placed next to one another (24 x 42 m), while block 1 was separated into two sub-blocks (each 12 x 42 m) (Figure S1). In mid-March 2019, pre-crops were chemically terminated and incorporated into the soil. To further boost the contrasts between the four *M. chitwoodi* densities (RKN densities), pre-crops (Table 1) were re-sown on May 7^th^ and mowed on July 16^th^. Right before the sowing of the cover crops, pre-crop stubble was milled and incorporated into the topsoil. To measure the soil nutritional status after pre-crops, 500 g of bulk soil was collected per strip and then pooled in four composite samples, one per pre-crop. Chemical analyses (Eurofins Agro, Wageningen, NL) gave the following results: C/N ratio = 18 - 21, total nitrogen: 3,750 – 4,280 kg ha^-1^, total phosphorus (P): 1,125 – 1,240 kg ha^-1^, with plant available P: 20.7 - 29.1 kg ha^-1^; total potassium (K): 110 – 270 kg ha^-1^, with plant available K: 110 – 155 kg ha^-1^.

**Table 1.**
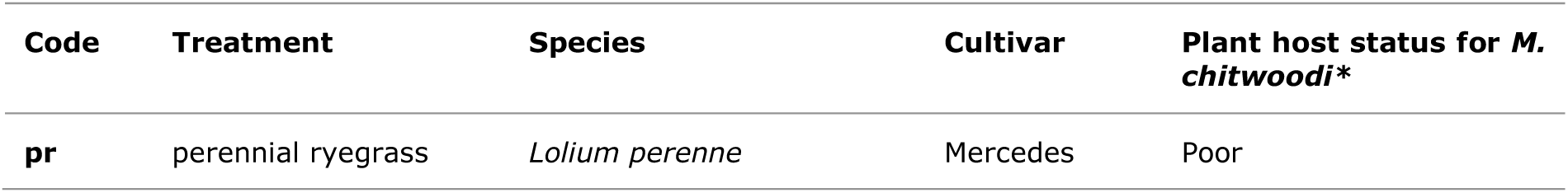
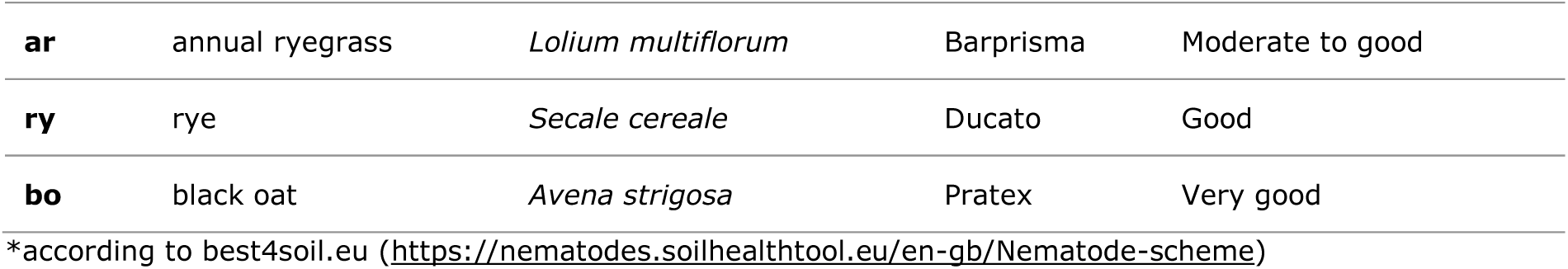
Pre-crops grown in the experimental field between August 2018 and July 2019 to generate distinct *Meloidogyne chitwoodi* levels.

### 2.2 Selection of cover crop species

Cover crops were sown on August 7^th^ (2019). Per strip (6 x 42 m), 11 plots (6 x 3 m, ≈ 0.9 m spacing between plots) were defined to accommodate six cover crop monocultures, four cover crop mixtures and one unplanted control (fallow) (Table 2). Every cover crop treatment and the fallow control were represented in each strip. Cover crop treatments were sown in the same order per block and in randomized order between blocks (Suppl. Fig. S1). Cover crops were mowed on December 2^nd^ (2019), and the plant residues were incorporated into the topsoil with a rotary tiller.

**Table 2.**
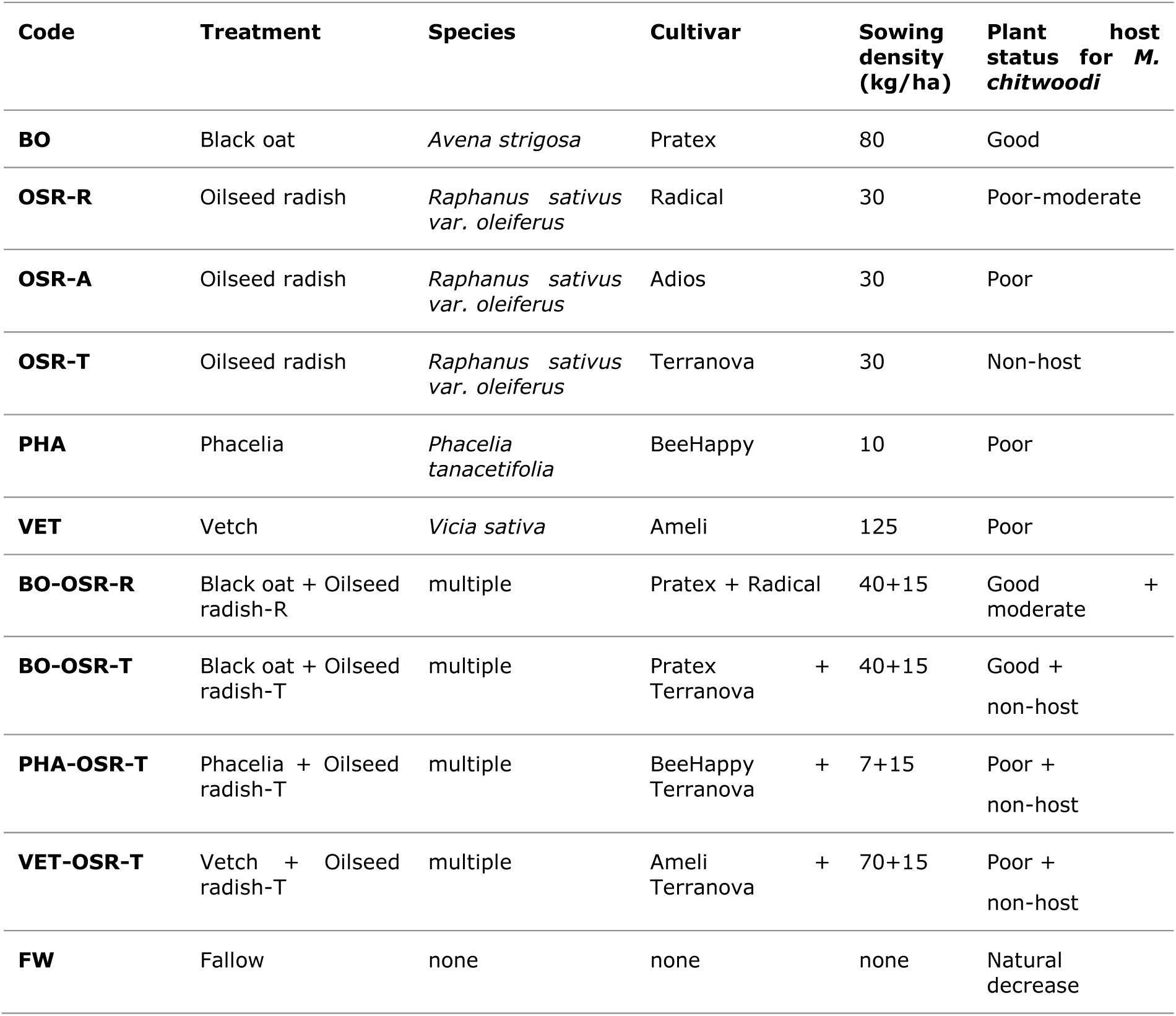
Details of the cover crop species and cultivars used in this study, including the origin of seeds, sowing density and expected host status for *Meloidogyne chitwoodi*.

### 2.3 Soil sampling for determination of *M. chitwoodi* densities

To determine the effect of cover crops on *M. chitwoodi* densities, bulk soil samples were collected just before sowing the cover crops (August 5^th^, 2019; T0_nem_) and on the day of cover crop termination (December 2^nd^, 2019; T1_nem_) (Figure 1). One litre of topsoil soil was collected with augers (∅ 12 mm, core length 25 cm) from the central area (1.5 × 2.7 m) of each plot. The soil sampled from each plot was carefully mixed, after which a subsample of 100 mL (≈ 120 g) was taken to determine the nematode densities. Soil subsamples were rinsed over 180 μm sieves. The organic material (> 180 μm) remaining on the sieve was incubated for four weeks at 20°C to allow the eggs present in the sample to mature and hatch (= ‘incubation fraction’). The soil suspension that passed the sieves (= ‘mineral fraction’) was extracted with an Oosterbrink funnel, and concentrated on three stacked 45 μm sieves. The material collected on these sieves was incubated on a filter for three days at 20 °C. The resulting nematode suspension was concentrated in 100 mL of tap water. *M. chitwoodi* individuals were counted under a microscope (Leica DMi8, 40x or 400x magnification) for two 10 mL subsamples from both the mineral and incubation fractions. When < 100 *M. chitwoodi* juveniles were found in a 10 ml subsample, the number of *M. chitwoodi* nematodes in the residual suspensions was counted as well. Soil sampling and nematode counting were conducted at the facilities of WUR-FC in Lelystad (NL).

**Figure 1.**
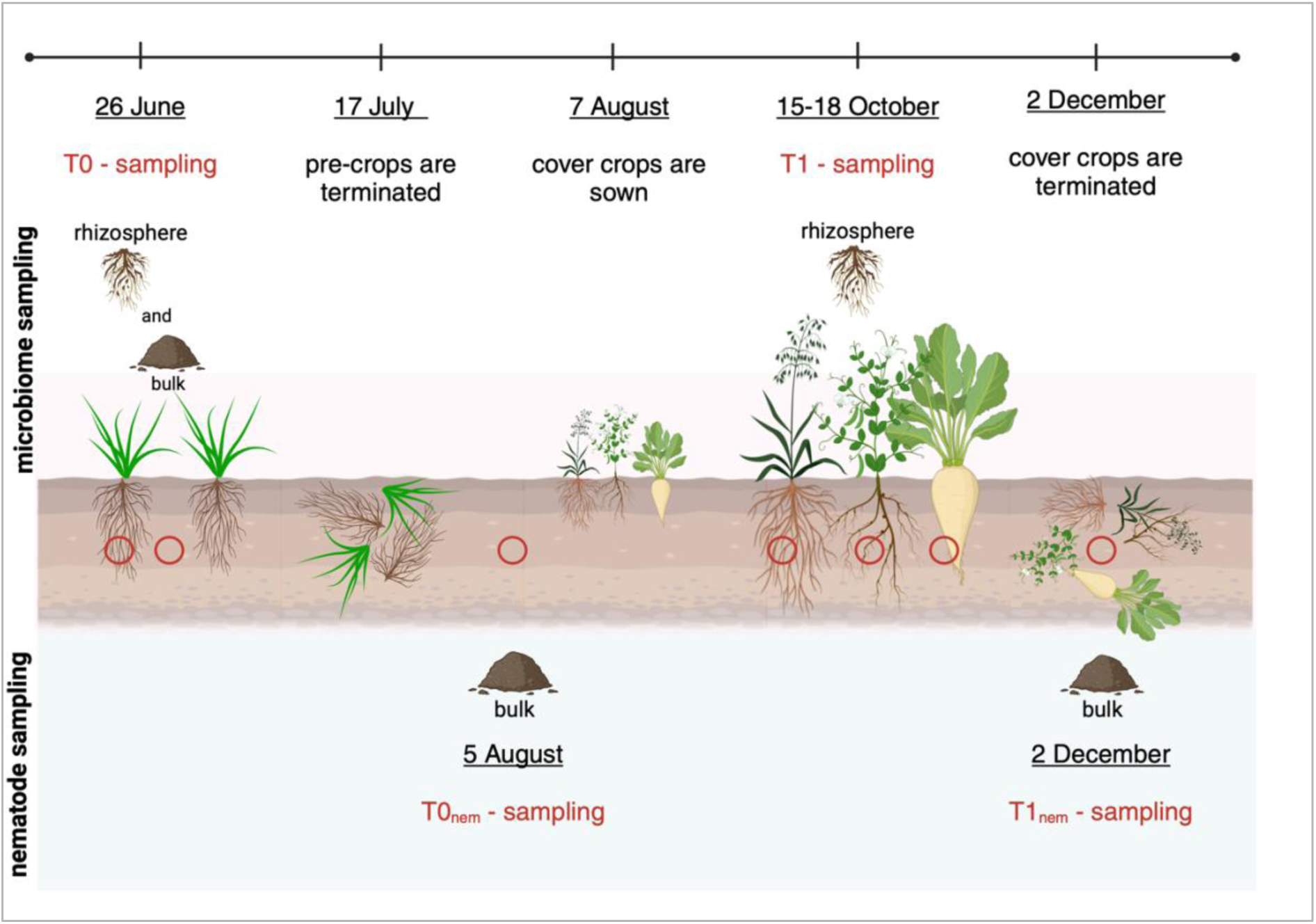
Timeline of the experimental field. To determine the initial microbial composition, both bulk and rhizosphere soil were sampled on June 26^th^ during pre-crop growth (‘T0 - sampling’). After pre-crop termination (July 17^th^) soil was collected to measure the initial *M*. *chitwoodi* densities (‘T0nem – sampling’). Cover crops were sown in early August, and rhizosphere soil was sampled from cover crops while still in vegetative growth stage (October 15-18^th^) to study the rhizosphere microbiome composition of each cover crop treatment (‘T1 – sampling’). Cover crops were terminated on December 2^nd^, and on the same day soil was collected to assess the final *M*. *chitwoodi* densities in the plots after cover crops (‘T1nem - sampling’).

### 2.4 Soil sampling for microbiome analyses

The first soil sampling (‘T0’) (Figure 1) was conducted on June 26^th^ (2019), during pre-crops growth. For each strip, bulk soil and rhizosphere soil were collected. Bulk soil was sampled by taking soil cores in between plants with an auger (∅ 12 mm, core length 20 cm). Each strip was randomly sampled by collecting 20 cores while avoiding the strip edges. Cores were thoroughly mixed and subsequently sieved (using a 2 mm mesh sieve). In total 12 bulk soil samples (4 pre-crops treatments x 3 replicates) were collected. Next to this, four randomly selected pre-crop plants were carefully extracted from each strip and taken to a nearby WUR-FC lab facility. Roots were mildly shaken to remove non-rhizosphere soil. Subsequently, rhizosphere soil was collected by gently brushing off soil adhering to the roots. Rhizosphere samples from four plants from the same strip were lumped and mixed. Hence, in total 12 rhizosphere samples (4 kinds of pre-crops x 3 replicates) were collected.

Cover crop rhizosphere soil was collected on the 15^th^ and the 18^th^ of October 2019 (‘T1’) (Figure 1). Rhizosphere soil was collected from the roots of four plants per plot. In the case of cover crop mixtures, the same amount of rhizosphere soil was collected from four couples of allospecific neighbouring plants. Three soil cores were collected from each of the fallow plots with an auger (diameter 12 mm, core length 20 cm). Soil and plants were collected in the central part of the plots (1.5 x 3 m) to avoid a border effect. Hence, per strip 11 samples were collected at T1, and this should have resulted in 132 rhizosphere soil samples. However, two samples were lost during the sampling process, so effectively 130 samples were analysed. For all rhizosphere and bulk soil samples, subsamples of 10 g were taken, snap-frozen in liquid nitrogen and transported on dry ice to the Laboratory of Nematology (WUR).

### 2.5 Illumina NovaSeq sequencing of 16s and 18S rDNA and rRNA

For all 10 g subsamples, an aliquot of 2 g was taken to isolate total DNA and RNA using an in-house developed, phenol-chloroform-based extraction protocol (Harkes et al., 2019). cDNA was synthesized from the isolated RNA using the Maxima First Strand cDNA Synthesis Kit for RT-PCR (Fermentas, Thermo Fisher Scientific Inc., USA) following the manufacturer’s instructions. Library amplification was performed on DNA and cDNA extracts with the following primer combinations: 515F/806R (Caporaso et al., 2012) targeting the V4 region of 16S (bacteria) and gITS7/ITS4ngs (Tedersoo et al., 2018) targeting the ITS2 region of fungi. DNA and cDNA samples were diluted to 1 ng μl^-1^ and 0.1 ng μl^-1^ respectively. Three μl of the diluted samples were mixed with 10 µl of IQ Supermix (Bio-Rad Laboratories, Inc.), 5 µl of Milli-Q and 1 µl of 5μM primers. For the first PCR reaction, the following temperature profile was used: 3 min. at 95°C for initial denaturation, followed by 35 cycles of 10s at 95°C, 30s at 56°C, 30s at 72°C and a final extension time of 5 min. at 72°C. After the first amplification, PCR amplicons were diluted 40-fold with Milli-Q water. Two μl of the diluted product was combined with 5 μl of Phire Hot Start II PCR Master Mix (ThermoFisher Scientific), 2 μl of Milli-Q water and 0.5 μl of forward and reverse primers (5 μM). These primers included the Illumina sequencing adaptors and index sequences for sample multiplexing. For the second PCR reaction, the following temperature profile was used: 3 min at 98°C for initial denaturation, followed by 15 cycles of 10 s at 98°C, 30 s at 60°C, 30 s at 72°C and a final extension time of 5 min at 72°C. Control samples including Milli-Q water only were taken along in the library preparation. Gel electrophoresis was used to check for correct amplicon size and purity for a random selection of PCRs products. Amplicons were pooled in one library and size selection and clean-up were carried out using AMPure XP Reagents (Beckman Coulter, Inc.). Finally, the library was sequenced using a standard Illumina NovaSeq SP2 (2×250bp) protocol (Illumina, San Diego, CA) and demultiplexed at Useq (Utrecht, The Netherlands). Sequences are available online in the NCBI SRA (Sequence Read Archive) database under BioProject number PRJNA973547.

### 2.6 Pre-processing of raw sequence data

Demultiplexed reads were sorted into the two organismal groups based on their locus-specific primer sequences. Forward and reverse reads were paired (merged on overlapping sequences) and clustered into amplicon sequence variants (ASVs) using the DADA2 pipeline (Callahan et al., 2016). For the ITS dataset, we applied the following filtering parameters: maximum expected error (maxEE) of 2 for both forward and reverse reads, truncation quality (truncQ) of 2 and no truncation (truncLen) of the amplicons, as advised for ITS reads. Taxonomic assignment for the ITS dataset was carried out using the IdTaxa method from DECIPHER (Wright, 2016), and the UNITE_v2021 database. For the bacterial dataset, we employed a maxEE of 2 for both forward and reverse reads, a truncQ of 2, and a truncation length (truncLen) of 230 for both forward and reverse reads. Taxonomic assignment for the bacterial dataset was performed using the default DADA2 method against the database SILVA_SSU_v138. As the quality assumption of the base call is differs between NovaSeq as compared to MiSeq, we applied a non-default method to estimate the error rate by using the DADA2 *errorEstimationFunction* parameter in the *learnErrors* function (https://gist.github.com/Jorisvansteenbrugge/a4f26030a047af6197b37f410f189fd4). Phyloseq objects (McMurdie and Holmes, 2013) were created for the bacterial and the fungal ITS datasets by merging the ASV tables, taxonomy tables and phylogenetic trees resulting from the DADA2 pipeline and the metadata tables. The decontam package (Davis et al., 2018) with default settings was used to compensate for bacterial or fungal contaminants in the laboratory. For this, the ASV composition of the control samples (MilliQ water, standard sample handling) was used. No fungal signals were detected in these water controls.

We discarded ASVs that were not annotated at the highest taxonomic level (phylum), singletons (accounting for 1.8% and 1% of the total bacterial and fungal ASVs, respectively) and ASVs belonging to non-target organismal groups. After this step, only annotated bacterial ASVs were included in the bacterial phyloseq and fungal ASVs in the fungal-ITS dataset. Furthermore, we filtered out rare ASVs (present in one sample only) and ASVs represented by less than 10 reads. Samples with < 10,000 bacterial and < 1,000 fungal reads were discarded (true for 1 DNA (bacteria) and 1 RNA sample (fungi)).

For bacteria, 29,890,835 reads were generated at the DNA level which clustered into 15,489 ASVs. The median number of reads and ASVs per sample were 193,687 and 1,675 in 153 samples (one DNA sample had < 10,000 reads). At the RNA level, 28,439,842 reads and 17,302 ASVs were produced from 154 samples, with a median read number of 184,569, and a median ASV number of 1,546.

From the fungal dataset, based on ITS sequencing, 22,013,785 DNA reads were retained after pre-processing, which clustered into 2,850 ASVs. The median number of reads and ASVs per sample were respectively 127,284 and 201 (154 samples). At the RNA level, 15,087,455 reads were employed that clustered into 2,011 ASVs. The median number of reads per sample was 93,148, while the median number of ASVs was 177 for 153 samples (one RNA sample had < 1,000 reads). The filtered bacterial and fungal phyloseq objects were used as input for statistical analyses.

### 2.7 Selection of putative nematode antagonists towards *M. chitwoodi*

Based on the reviews by Li et al. (2015) and Topalovic et al. (2020), we made an overview of bacterial and fungal genera harbouring at least one species that has been described as an antagonist of plant-parasitic nematodes (Suppl. Table S1). We used this overview to identify genera harbouring nematode antagonists in our datasets. We refer to the shortlisted genera as putative nematode antagonists throughout the study for simplicity.

### 2.8 Statistical analyses

All statistical analyses were performed using R v.4.2.2 (R Core Team, 2021) with the aid of relevant packages, including phyloseq v.1.42.0 (McMurdie and Holmes, 2013) and vegan v.2.6-4 (Oksanen et al., 2013).

#### 2.8.1 Effect of pre-crops on RKN densities

Generalised linear mixed models with negative binomial error distributions (glmm-nb) were used to assess the effects of pre-crops (at T0_nem_) and cover crop treatments (at T1_nem_) on *M. chitwoodi* densities. Raw nematode counts were used as response variables in the models, with block as a random factor to account for the positional effect of the field. Analysing the effect of pre-crops on the T0_nem_ counts, we determined the initial population density levels of *M. chitwoodi*. These levels will be referred to as ‘RKN densities’. Analysing the effect of cover crop treatments and RKN densities on the T1_nem,_ an interaction factor (cover crop treatments * RKN density) was added to the model.

#### 2.8.2 Effects of pre-crops, cover crop, and RKN densities on soil microbiome

The effects of pre-crops, cover crop treatments and the distinct *M. chitwoodi* levels on the total and active bacterial and fungal community structure was assessed by Permutational analysis of variance (PERMANOVA – *adonis2* function in vegan) and Principal Coordinate Analysis (PCoA – *plot_ordination* function in phyloseq). For both tests, ASVs were normalised through Cumulative Sum Scaling (CSS) transformation (Paulson et al., 2013), and the distance matrix was calculated using the Bray-Curtis dissimilarity index. As PERMANOVA tests each term in sequential order, the positional effect of blocks was accounted for by introducing the term ‘block’ as the first term in the PERMANOVA. The same tests, PERMANOVA and PCoA, were used to assess the community structure of the selection of putative nematode-antagonistic microbial genera. Cover crop treatments were compared with pairwise PERMANOVAs (*pairwise.perm.manova* function, RVAideMemoire v.0.9 (Hervé and Hervé, 2020)) carried out based on Bray-Curtis multivariate distances with Benjamini-Hochberg correction for multiple testing and 999 permutations.

#### 2.8.3 Effects of cover crop treatments on the diversity of putative nematode antagonists

Alpha diversity indices describing richness (Observed index) and diversity (Shannon index) were calculated on the selection of putative nematode antagonists from the rarefied bacterial and fungal ASVs table using the phyloseq package. Rarefying was conducted to the minimum library size without replacement. The smallest library size between DNA and RNA datasets was used for bacteria and fungi (RNA had in both cases the smallest library size). Differences in alpha diversity scores among pre-crops were tested with linear models, using the logarithm of the Observed and Shannon index as the response variables. The effects of cover crop treatments were tested with the Kruskal-Wallis rank sum test (KW), after rejecting the normality of data and residuals from linear models implemented with raw or log-transformed data. As KW can only test one factor at the time, the effect of pre-crops and the positional effect (= block effect) on the T1 samples were also tested separately and reported if significant. When the KW statistical test was significant, post-hoc tests were conducted with Dunn’s test and Holm p-value adjustment for multiple testing with ggstatsplot v 0.10.0 (Patil, 2021) which was also used to generate the plots.

Rarefied ASV tables were also used to calculate the relative abundance of putative nematode-antagonistic genera per pre-crop and cover crop treatment at DNA and RNA levels which were potted in barplots.

#### 2.8.4. Effects of cover crops and nematode densities on the abundance putative nematode antagonists

Glmm-nb models were used asses if the RKN densities and cover crops related to the abundance of putative bacterial and fungal nematode antagonists (Zhang and Yi, 2020). ASVs of putative nematode antagonists were aggregated to genus level, and raw DNA and RNA read counts of each genus were used as a response variable in the mixed models. Only antagonists with a prevalence of > 25% (present in > 3 and > 33 samples at T0 and T1, respectively) were tested with models. To account for differences in the sequencing depth, the logarithm of the total read counts per sample was added as an offset term. In addition, the block term was introduced as a random effect in the models to account for the positional effect. When assessing the effect of cover crops and RKN densities on the read counts of the antagonists, these two terms were first added as an interaction effect (cover crop treatments * RKN densities). In the absence of a significant interaction term, the individual terms were reciprocally included as both fixed and random effects in two models. In these models, the fallow samples (bulk soil) were excluded and only rhizospheric samples from cover crops were tested. Glmm-nb were implemented in glmmTMB v. 1.1.6 (Brooks et al., 2017) and the fit of each model and zero inflation were tested with DHARMa v. 0.4.6 (Hartig, 2020). Zero-inflated negative binomial models were used when zero inflation was significant. Pairwise comparisons of estimated marginal means were assessed with emmeans (v. 1.8.5 (Lenth, 2022) and multcomp v.1.4-23 (Hothorn et al., 2008) and p-values were corrected with Benjamini-Hochberg procedure. To present the results of the models and show significant comparisons, box plots were built on the non-rarefied ASVs for those genera tested in the models (provided as supplementary material). All tests were considered statistically significant at P < 0.05.

## 3. Results

### 3.1 Effects of pre-crops on *M. chitwoodi* and soil microbiome

Growing four grass species with distinct host status for root-knot nematode *M. chitwoodi* significantly affected the densities of this plant parasite (Χ^2^_3,132_ =156.57, P < 0.001; R^2^ = 63.47%, Fig. 2A) at T0_nem_. Growing black oat as a pre-crop, a very good host for *M. chitwoodi*, resulted in the highest nematode densities (median_bo_ = 5,650 individuals 100 g soil^-1^). Growing rye (ry) and annual ryegrass (ar) gave rise to intermediate *M. chitwoodi* densities with medians of 938 and 401 individuals 100 g soil^-1^, respectively. Exposure to perennial ryegrass (pr) resulted in establishment of the lowest *M. chitwoodi* levels (median_pr_ = 1 individual 100 g soil^-1^). As the rye (ry) and annual ryegrass (ar) treatments did not result in significantly distinct RKN levels (Fig. 2A), the four treatments with grass species resulted in the generation of three distinct RKN levels (‘high’, ‘medium’ and ‘low’ RKN in Fig. 2B).

**Figure 2.**
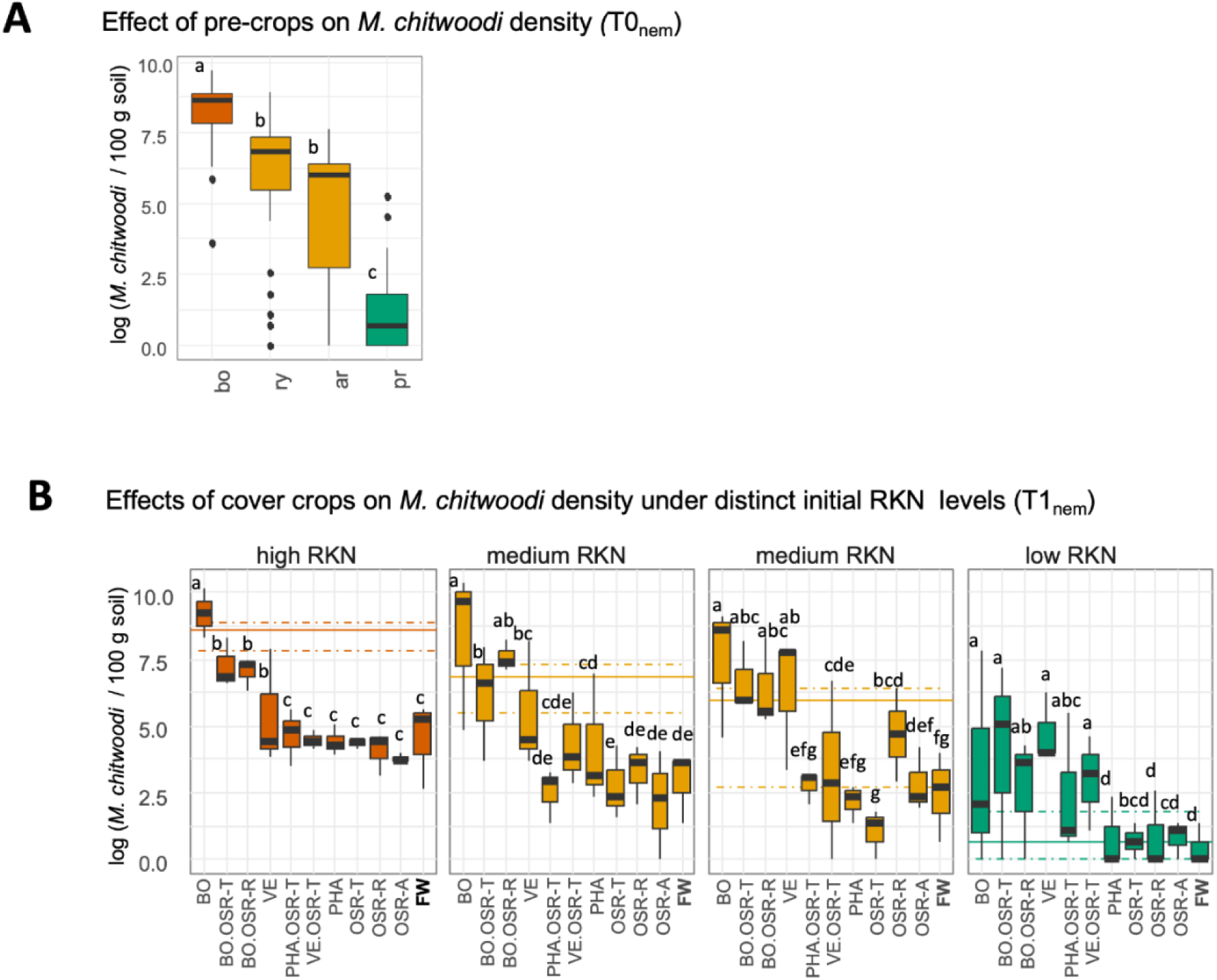
*M. chitwoodi* densities after pre-crops (T0_nem_) and after the cover crop treatments (T1_nem_). A. Four pre-crops were grown: black oat (bo), rye (ry), annual ryegrass (ar) and perennial ryegrass (pr). Different letters above box plots indicate statistically significant differences among the pre-crop treatments. B. To visualize the impact of cover crops in plots with distinct RKN levels, initial RKN levels are provided in each of the four graphs. Solid lines and dotted lines indicate the median and respectively the 0.25 and 0.75 quantiles of initial *M. chitwoodi* density counts for each pre-crop (T0_nem_). Cover crop treatments are abbreviated and further explained in Table 2. Box plots show the impact of cover crops on *M. chitwoodi* for each of pre-crop treatments.

To pinpoint the impact of the pre-crops on the bacterial and fungal communities both rhizosphere and bulk soil were sampled at T0 (Fig. 1, Table 1). At T0, similar bacterial and fungal community structures were found at both DNA and RNA level for the four pre-crop treatments irrespective whether rhizosphere or bulk soil was sampled (PERMANOVA after removing the variation due to block effect, P > 0.05, Suppl. Table S2 A, Suppl. Fig. S2). As the four poaceous pre-crops did not result in distinct shifts in the bacterial and fungal communities, but generated three distinct RKN levels, plots were pooled on the basis of their RKN densities in the following microbiome analyses (Suppl. Table S2 A)

### 3.2 Effects of cover crop treatments on *M. chitwoodi*

At the time of cover crop termination (T1_nem_), a strong interaction was observed between the pre-crop treatments and the subsequent impact of the cover crops (Χ^2^_30,132_ = 65.01, P < 0.001, R^2^ = 80.33%). Therefore, the effect of each cover crop treatment on *M. chitwoodi* was assessed separately for each of the four pre-crops (Fig. 2B). For all pre-crop treatments, cover crop treatments including black oat (*viz.* monoculture BO and the mixtures BO-OSR-T, BO-OSR-R, see Table 2) showed the highest *M. chitwoodi* densities (range: median_treatment with BO_= 7 (after perennial ryegrass) - 15,970 (after rye) individuals 100 g soil^-1^, Fig. 2B). Plots exposed to fallow (FW), oilseed radish monocultures (OSR-T, - R and -A), and phacelia (PHA) typically showed the lowest *M. chitwoodi* densities (Fig. 2B). This is further illustrated by the impact of oilseed radish cv. Terranova resulting in median densities between 1 (= lowest after perennial ryegrass) and 83 (= highest after black oat) *M. chitwoodi* individuals 100 g soil^-1^.

### 3.3 Effect of cover crop treatments on the overall microbiome community compositions

At T1, rhizosphere soil from all ten cover crop treatments and bulk soil from the fallow plots were collected, and microbiome analyses revealed that all cover crop treatments significantly affected the resident (DNA) and the active (RNA) fractions of the bacterial and fungal communities (Fig. 3) (PERMANOVA, effects cover crops: R^2^_bact-DNA_ = 22%, R^2^_bact-RNA_ = 25%; R^2^_fun-DNA_ = 27%, and R^2^_fun-RNA_ = 21%, for all P < 0.001). Initial RKN densities had no significant effect on the bacterial DNA and RNA communities, and only explained 2% of variation for the fungal communities (both at DNA and RNA level) (P < 0.01) (Suppl. Table S2 B). The interaction effect of cover crop treatments and RKN densities was not significant (PERMANOVA, P > 0.05) (Suppl. Table S2 B).

**Figure 3.**
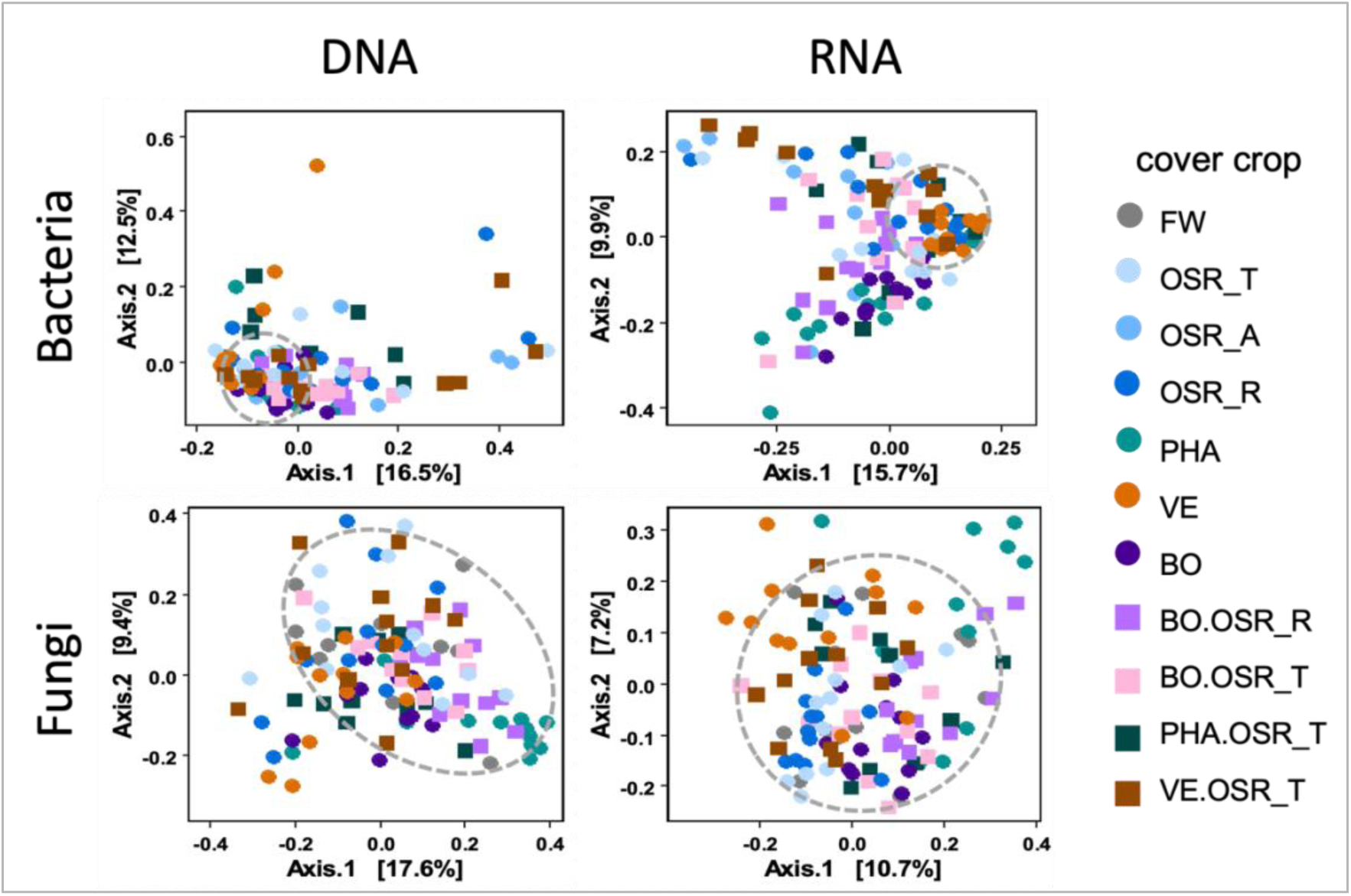
Community composition of the rhizospheres of six cover crop monocultures, four mixtures and fallow bulk soil at T1. Colours indicate cover crop treatments; shapes are used to discriminate between monocultures (spheres) and mixtures (squares). Principal Coordinate Analysis (PCoA) of CSS-normalised ASVs based on Bray-Curtis-based dissimilarity matrices show that microbial communities are separated along the axes as cover crop treatment accounted for 10.7-17.6% of variation along the principal PCoA axis (PERMANOVA, p ≤ .001). Dashed circles are used to group microbial communities in fallow controls (grey spheres).

Pairwise comparisons of cover crop treatments (Suppl. Table S2C) revealed significant differences between all rhizosphere communities and the bulk communities in the fallow plots (P < 0.01). Furthermore, no differences were observed in rhizobiome composition between the oilseed radish cultivars Adios, Radical and Terranova (P > 0.05).

In most cases, the microbiome of cover crop mixtures that included oilseed radish Terranova did not significantly differ from the Terranova monoculture (P > 0.05). On the other hand, the microbiome composition of cover crop mixtures such as BO-OSR-T, PHA-OSR-T and VET-OSR-T deviated from the microbiomes in the rhizospheres of monocultures of black oat, phacelia and vetch. These results revealed that in case of cover crop mixtures, oilseed radish cv. Terranova had a dominant effect on the composition of the rhizosphere microbiome (P < 0.01). This dominance was not a characteristic for all oilseed radish cultivars since the microbiome of the black oat - oilseed radish cv. Radical mixture (BO-OSR-R) deviated significantly from the microbiomes associated with each of the constituents (P < 0.01).

### 3.4 Diversity of local nematode antagonists detected in the experimental field

Based on two authoritative reviews (Li et al. 2015, Topalovic et al. 2020), we classified eight bacterial and 26 fungal genera as nematode antagonists *sensu lato* (Suppl. Table S1). Also, fungal genera harbouring species with an endophytic lifestyle strengthening plant defence responses against plant-parasitic nematodes were included in this list (Suppl. Table S1). Especially for fungi, the afore-mentioned number is only a general approximation as the systematics of some of these genera is still volatile. It is noted that genera such as *Arthrobotrys* exclusively harbour nematode antagonists (Zhang et al., 2022), while, for example, within the speciose genus *Fusarium,* nematode antagonists constitute only a small minority (Benitez-Malvido et al., 2021). Within the 130 T1 rhizospheric soil samples analysed from the Vredepeel experimental field, five bacterial and 14 fungal genera comprising putative nematode antagonists were detected, covering 63% and 54%, respectively, of the overall described antagonist diversity.

Five out of the eight bacterial genera harbouring nematode putative antagonists were prevalent and active in almost all samples (98-100% at the DNA level; 97-99% detected at the RNA level). Within four of the five genera with putative nematode antagonists, the list of species found in the experimental field included taxa that were shown in literature to act as antagonists (*e.g.*, *Pseudomonas putida* as a known nematode antagonist within the genus *Pseudomonas* (see also Table 3).

**Table 3.**
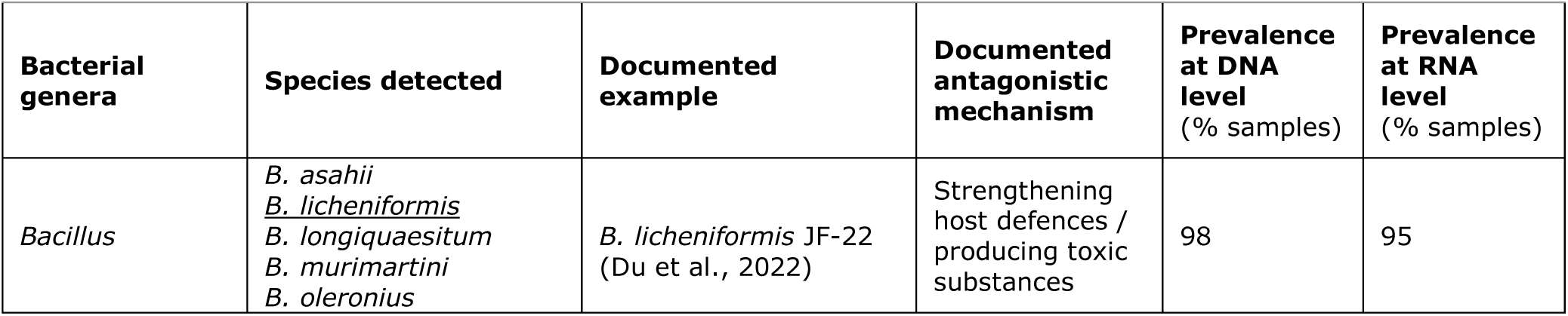
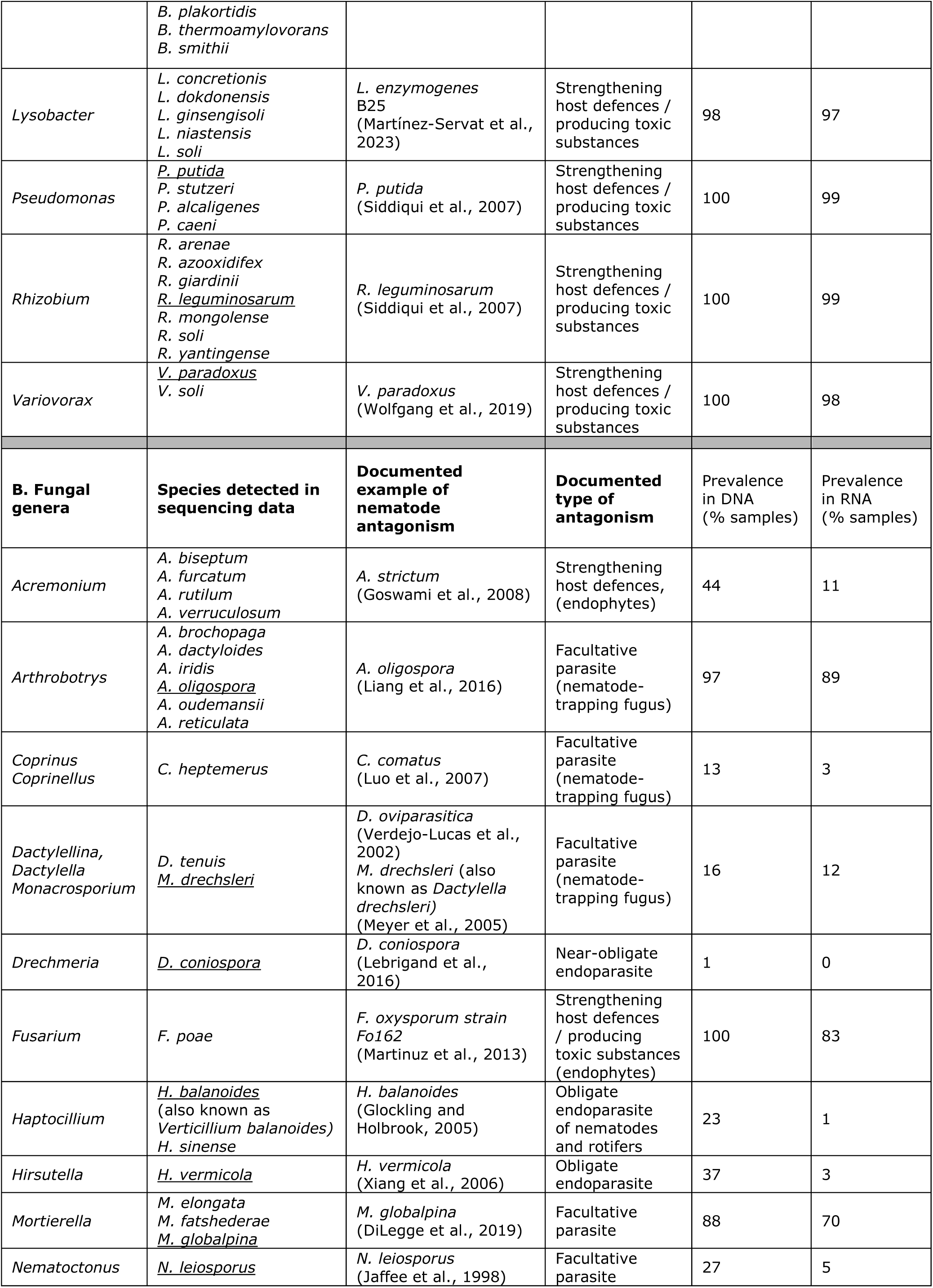
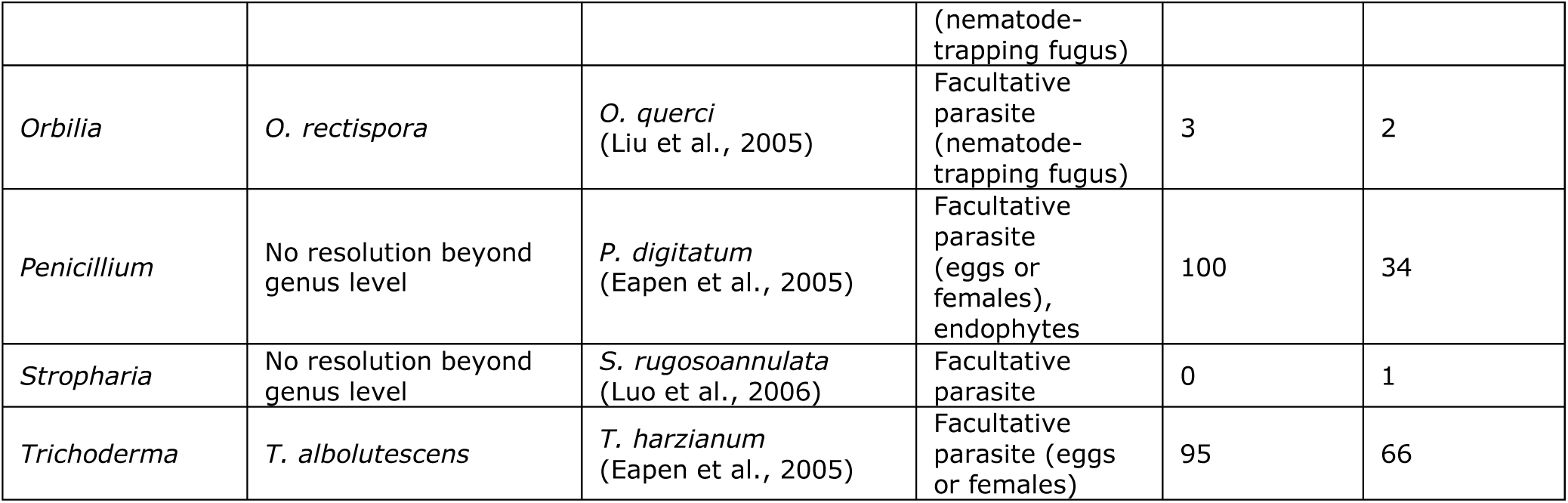
Overview of microbial genera found in the Vredepeel experimental field at T1 that are known to harbour nematode antagonists. ‘Species detected’ provides list of species found in the experimental field at the DNA level. Under ‘Documented example’ for each genus a single publication is presented that demonstrates nematode antagonism for a particular representative of that genus. In case the example species was found in the experimental field, the species name was underlined. Note that for many of the detected bacterial and fungal *species* it is unknown whether or not they interfere in plant-parasitic nematode – plant interactions. Prevalence at T1 (= % of the 118 rhizosphere samples harbouring members of a given genus) is determined at DNA (resident community) and at RNA level (active community).

Among the fourteen fungal genera harbouring putative nematode antagonists, large differences in distribution over the experimental field were observed (Table 3). Whereas members of the genus *Arthrobotrys* were present and active in around 90% of the samples, *Nematoctonus* was shown to be present in 27% of the samples while they were active in 5% of the samples only (Table 3). For several genera of putative nematode antagonists (*e.g.*, *Hirsutella*), we detected a species that is indeed known as a nematode antagonist (*e.g.*, *Hirsutella vermicola,* Xiang et al. (2006)). However, some fungal species identified within our dataset, *e.g.*, *Coprinus heptemerus,* could not be linked to nematode suppression.

### 3.5 Initial composition of the community of putative nematode antagonists under distinct RKN densities

At T0, we investigated whether the pre-crop treatments that gave rise to three distinct RKN densities had resulted in differences in putative nematode antagonist communities. No such an effect was detected as both richness (Observed) and diversity (Shannon) of the bacterial and fungal community did not differ among the three RKN densities (P > 0.05 for all combinations) (Suppl. Fig. Figure S3 B, D). Overall, the putative nematode antagonists’ community, as presented in Table 3 (and Suppl. Table S1) in the rhizosphere was uniformly distributed among the three RKN densities at T0 (PERMANOVA, P > 0.05 at DNA and at RNA level for bacteria and fungi, Suppl. Fig. S3 A, C, Suppl. Table S3).

**Figure 4.**
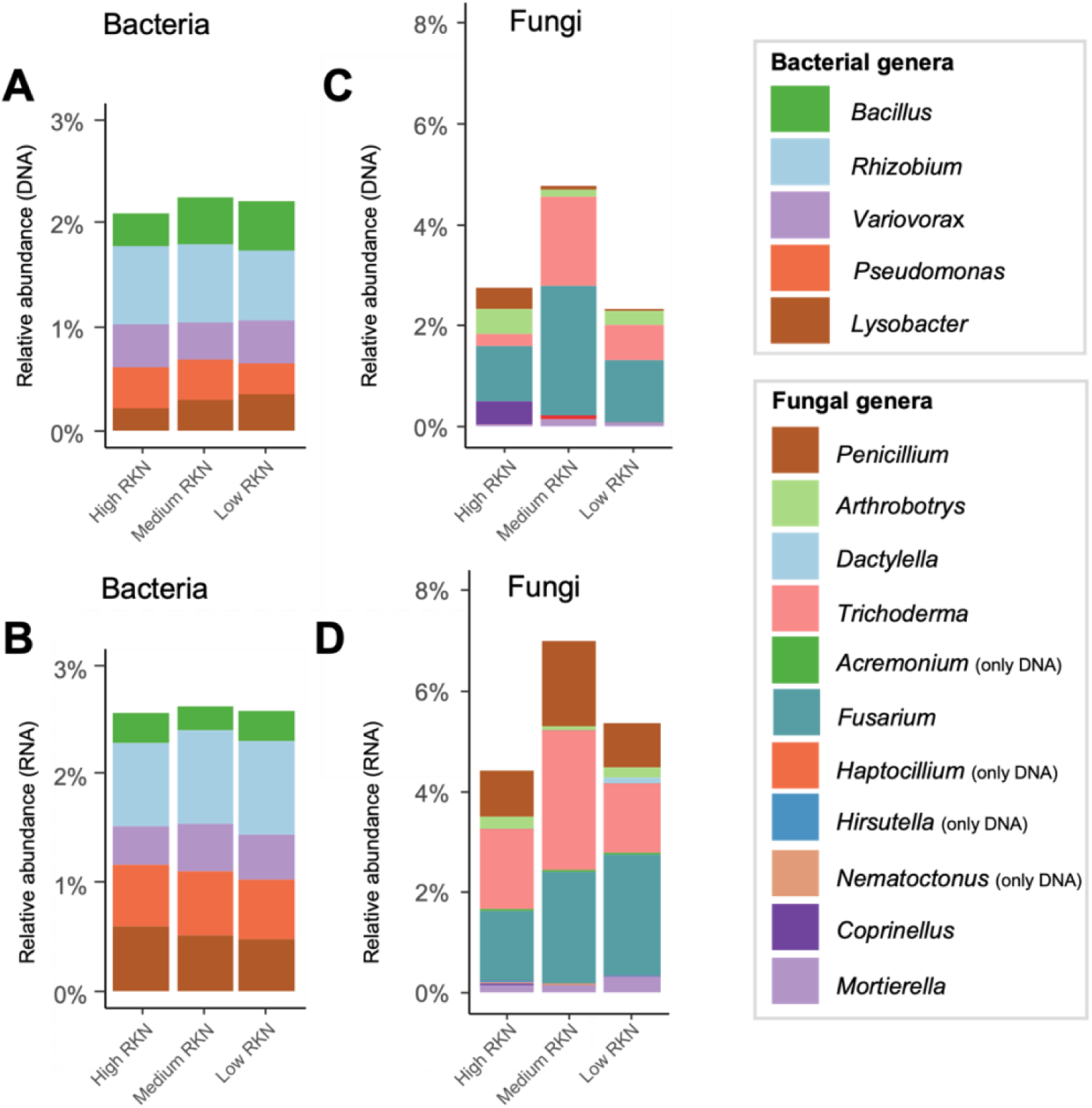
Relative abundances of microbial genera harbouring nematode antagonists in the rhizosphere of pre-crops at T0 under three distinct RKN densities (see also Fig 2A). Relative abundances were calculated as read counts for putative antagonists as fraction of the total read counts per sample * 100%. **A, B.** Resident (DNA) and active (RNA) bacterial genera that comprise nematode antagonists. **C, D.** Resident (DNA) and active (RNA) fungal genera that harbour nematode antagonists. In case genera were detected at RNA level only, this is specified behind the genus name.

### 3.6 Impact of cover crop treatments and RKN densities on the community of putative nematode antagonists

#### Bacterial putative nematode antagonists

At T1, cover crop treatments had a significant effect on the community structure of putative bacterial antagonists (PERMANOVA, P < 0.05, Suppl. Fig. S4, Suppl. Table S3 B) accounting for 31% to 19% of the microbiome variation at respectively DNA and RNA level (Suppl. Table S3 A). On the other hand, RKN densities did not affect the community structure of putative bacterial antagonists (PERMANOVA for RKN densities, P > 0.05). Diversity and richness of putative bacterial antagonists were significantly affected by cover crops both at DNA and RNA level (Table 4). In most cases, rhizosphere communities of putative nematode antagonists displayed a higher diversity and richness as compared to the bulk soil antagonist communities of the fallow (Suppl. Fig. S5 A, B).

**Table 4.**
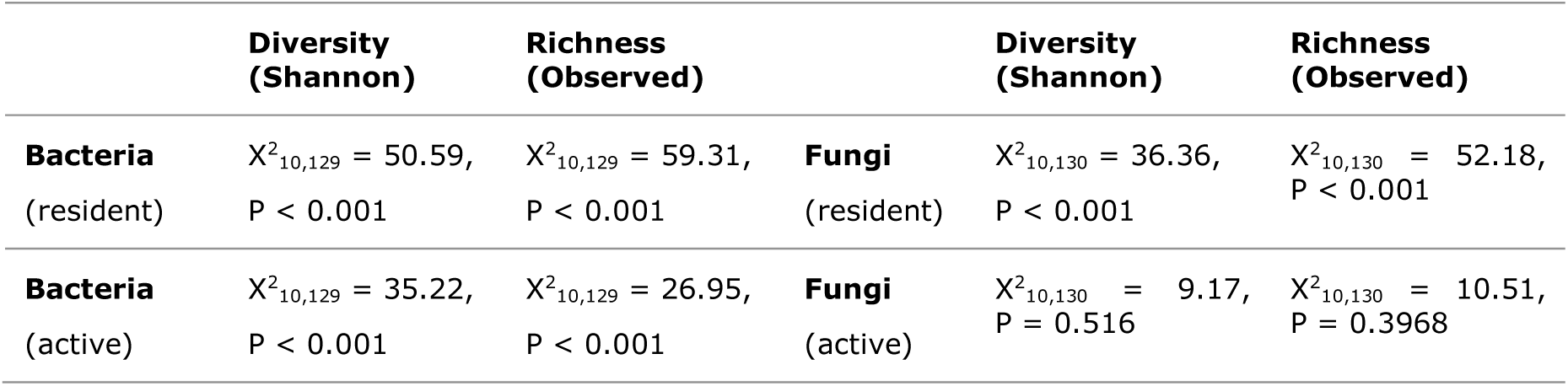
Effects of cover crop treatments on the diversity and richness of putative nematode-antagonistic genera as presented in Table 3. Diversity and richness indexes were calculated at the ASV level. P-values were generated with the Kruskal-Wallis rank sum test for non-normally distributed data.

In fallow plots at T1, genera harbouring nematode antagonists *sensu lato* accounted for 0.9% and 1.7% of the total bacterial community in bulk soil at DNA and RNA level respectively. Cover crop treatment resulted in an increase of the relative abundance putative nematode antagonists (Table 3) in the rhizosphere ranging from 2.7% in vetch, to 6.1% in oilseed radish cv. Adios at the DNA level (Fig. 5A), and from 1.6% in phacelia as well as black oat monoculture and mixtures to 2.5% in vetch at the RNA level (Fig. 5B).

**Figure 5.**
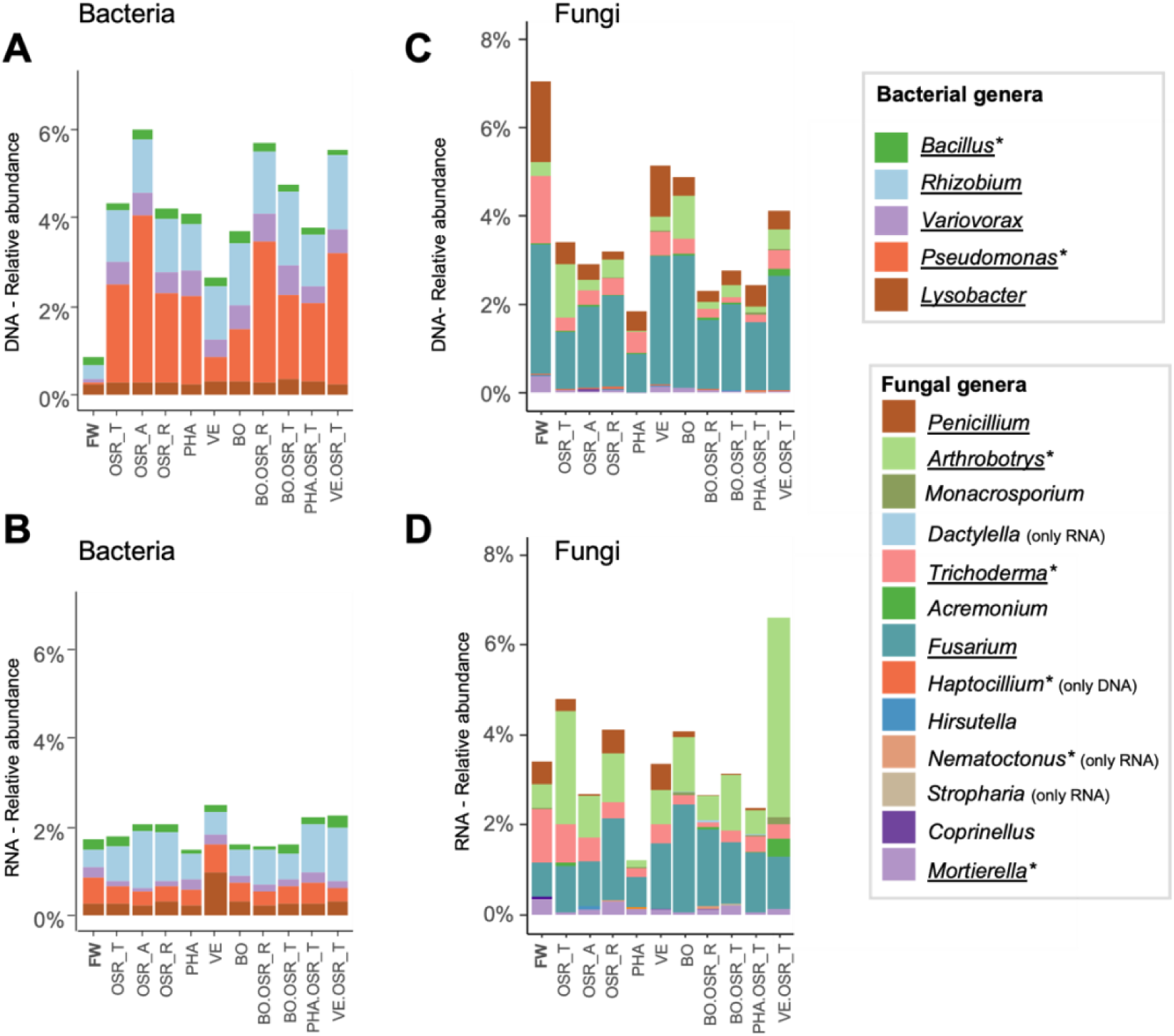
Relative abundances of microbial genera harbouring nematode antagonists in the rhizosphere of cover crops and the fallow control. Relative abundances were calculated as read counts for putative antagonists as fraction of the total read counts per sample * 100% at T1. Cover crop treatments are abbreviated, and further explained in Table 2. **A, B.** Resident (DNA) and active (RNA) bacterial genera comprising putative nematode antagonists. **C, D.** Resident (DNA) and active (RNA) fungal genera comprising putative nematode antagonists. In case genera were detected at DNA or RNA level only, this is specified behind the genus name. Genera that were detected at the DNA level in > 80% of the samples are underlined. Genera significantly responding to cover crop treatments and/or RKN densities are marked with asterisks (see also Table 5).

Model-based statistical analyses revealed that putative bacterial antagonists were stimulated by cover crops in a genus-specific manner (Figure 6). The abundance of *Pseudomonas* was influenced by the interaction between cover crop treatments and RKN densities. The vetch rhizosphere was characterized by the lowest *Pseudomonas* abundance (at DNA level) in combination with the highest *Pseudomonas* activity (at RNA level) (Figure 6, Suppl. Fig. S6). Monocultures of the oilseed radish cultivars Adios and Terranova, as well as all mixtures that included oilseed radish Terranova strongly promoted the resident *Pseudomonas* community (Figure 6).

**Figure 6.**
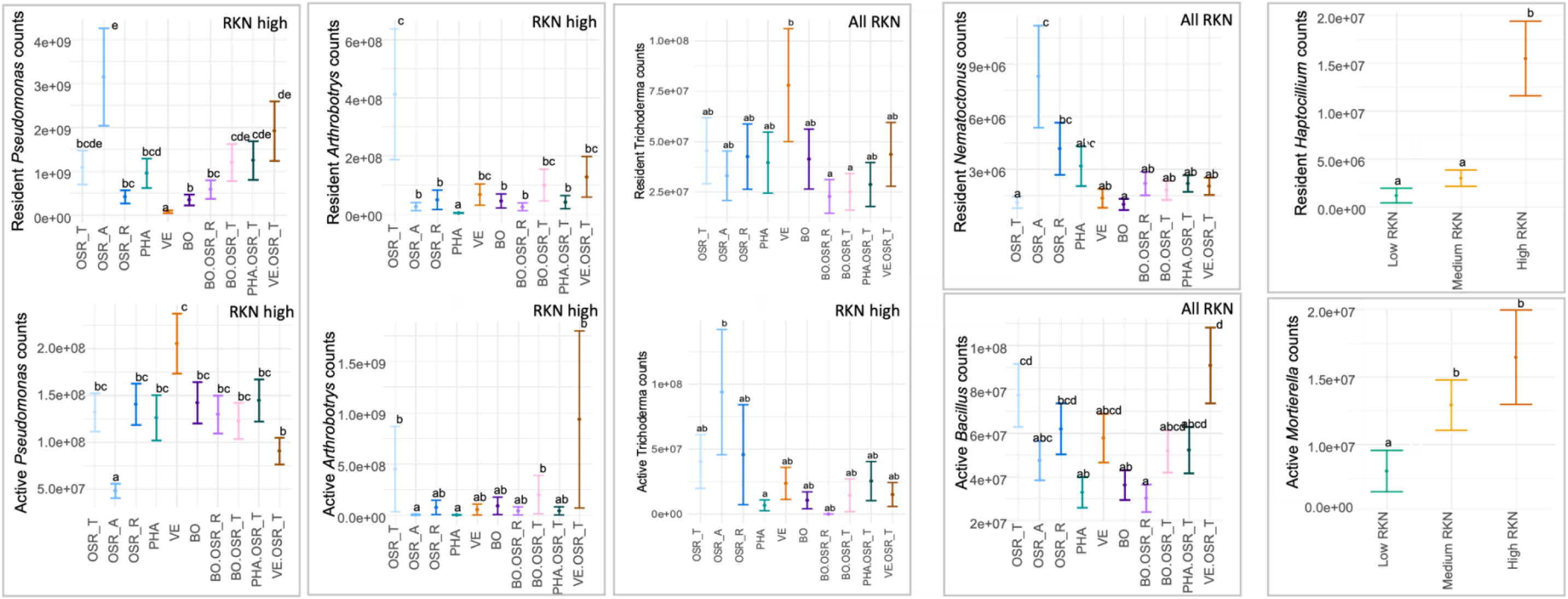
Effects of cover crops and *M. chitwoodi* levels on the abundance of bacterial or fungal genera comprising putative nematode antagonists at T1. Results of the negative binomial generalized linear mixed models are presented: estimated read number of putative genera per cover crop treatment and/or RKN density, and statistical significance between cover crop treatments or RKN densities assigned with post-hoc tests. A full overview of the effects of cover crops and RKN on nematode-antagonist harbouring microbial genera is presented in Suppl. Table S4. <u>Effects of cover crops</u>: When interaction between cover crop treatment and RKN density was significant, the effect of cover crop treatments was assessed per RKN density (Low, Medium and High). When the interaction was not significant, the effects were tested over all RKN densities (‘All RKN’). <u>Effects of *M. chitwoodi* levels</u>: only for two genera (*Haptocillium* and *Mortierella*) the effect of RKN density was significant. Different lowercase letters indicate significant differences (p < 0.05) in the read counts of putative nematode-antagonistic genera (calculated with post-hoc tests with Benjamini-Hochberg adjustment for multiple comparisons).

In contrast to *Pseudomonas,* representatives of the genus *Bacillus* were only affected at RNA level. *Bacillus* was activated in the presence of any cover crop mixture with oilseed radish cv. Terranova, as well as by monocultures of vetch, and oilseed radish cultivars Terranova and Radical, while at the DNA level no stimulation was observed.

#### Fungal putative nematode antagonists

Cover crop treatments accounted for ≈ 14% and 9% of the variation among fungal genera comprising nematode antagonists *sensu lato* at DNA and RNA level, respectively (P < 0.05, Suppl. Table S3 B, Suppl. Fig. S4). A significant effect of RKN densities on the putative nematode-antagonistic community was observed at DNA level (R^2^ = 2.8%, P < 0.05), but not at the RNA level. The interaction between cover crop treatments and RKN densities was non-significant (P > 0.05, Suppl. Table S3 B). Cover crop treatments had a significant impact on the diversity and richness (at the level of ASVs) of putative nematode antagonists at the DNA level, as most cover crops’ rhizosphere displayed higher richness and diversity of antagonists as compared to fallow (Table 4, Suppl. Fig. S5 C). At the RNA level, no significant effect of cover crop treatments on the diversity and richness of putative nematode-antagonistic genera was observed (Table 4).

In fallow plots at T1, fungal genera harbouring nematode antagonists *sensu lato* represented 7.1% at the DNA and 3.4% at the RNA level of the total community in bulk soil. At the DNA level cover crop treatment resulted in relative abundances of fungal genera comprising nematode antagonists between 1.8% and 5.1% in, respectively, phacelia and vetch (Fig. 5C). Focussing on active microbiota, putative nematode antagonists’ abundances ranged from 1.5% to 6.5% in phacelia and vetch-oilseed radish cv. Terranova mixture, respectively (Fig. 5 D). Three fungal genera were detected at RNA level only (Fig. 5 C, D). We hypothesize that representatives of these genera were present at a very low density, resulting in a DNA signal below the detection limit.

Model-based statistical analyses revealed that *Arthrobotrys* was significantly affected by the interaction of cover crops and RKN densities (interaction: X^2^_20,130_ = 50.10, P < 0.001, Suppl. Fig. S6). In plots with the highest RKN densities, this nematode-antagonistic genus was least abundant in phacelia and most abundant in oilseed radish cv. Terranova monoculture and its mixtures with black oat and vetch (at both DNA and RNA level, Figure 6, Suppl. Fig. S6). The abundance of *Trichoderma* spp. was influenced by cover crops only in high and medium RKN densities, while no difference in read counts was assessed at the low RKN density (Figure 6, and Suppl. Table S4). Under high RKN densities, *Trichoderma* showed increased activity, especially in plots treated with oilseed radish cv. Adios. The genus *Nematoctonus* (only tested at the DNA level, Figure 6) responded significantly to cover crops (X^2^ = 39.04, P < 0.001) and was most abundant in oilseed radish cv. Adios and least abundant in plots with black oat.

Two fungal genera harbouring obligate nematode parasites, *Hirsutella* and *Haptocillium* (both belonging to Ophiocordycipitaceae), were not affected by cover crops (tested at the DNA level only) (P > 0.05). *Haptocillium* abundance was affected by RKN density (X^2^_10,130_ = 22.72, P < 0.001), with the highest read counts associated with the highest RKN density (Table 5). The abundance of the genus *Mortierella* did not vary among cover crops, but its activity was significantly promoted upon exposure to medium and high RKN densities (Table 5, Suppl. Fig. S6).

It is concluded that both abundance and activity of microbial genera harbouring nematode antagonists can be stimulated by cover crops and by RKN density. It is noted that none of these two variables had a stimulatory or inhibitory effect on the conglomerate of bacterial or fungal genera comprising nematode antagonists as a whole.

## 4. Discussion

We investigated the impact of ten cover crop treatments on fungal and bacterial communities against a background of three density levels of the root-knot nematode (RKN) *M. chitwoodi* with a focus on microbial genera harbouring nematode antagonists. Bacterial and fungal taxa interacting with plant-parasitic nematodes have been extensively studied in the past, and this pre-existing knowledge allowed us to identify a substantial number of genera that are known to comprise putative nematode antagonists. In a conventionally-managed experimental field, five bacterial and 14 fungal genera were identified that comprise putative nematode antagonists. This represents 63% and 54% of, respectively, the bacterial and fungal genera that are described in two authoritative reviews on antagonists of plant-parasitic nematodes (Li et al. 2015, Topalovic et al. 2020).

We showed that both cover crop treatments and, to a lesser extent, RKN density impacted the abundance and/or activity of bacterial and fungal genera that comprise nematode antagonists. However, this selection of microbial genera responded distinctively to these treatments, and stimulation of the abundance of a taxon (at the DNA level) was not necessarily paralleled by an increase in the activity of the same taxon (at the RNA level) and *vice versa.* Overall, we observed a remarkably rich representation of bacterial and fungal genera associated with nematode antagonism in a common arable field, and these genera were shown to be more readily stimulated by cover crops than by putative prey (*M. chitwoodi*) densities.

### Nematode antagonists are abundant and diverse in a conventionally managed experimental field

Cohabitation of nematodes and soil microbes for hundreds of millions of years triggered the evolution of nematode antagonists. Among fungi, nematophagous lifestyles diverged from saprophytism about 419 MYA onwards (Yang et al., 2012). Two recent overviews by Li et al. (2015) and Topalovic et al. (2020) suggest that in at least eight bacterial and 26 fungal genera antagonism against plant-parasitic nematodes has evolved. Assuming that these numbers are in the correct order of magnitude, the ample representation of nematode antagonist-harbouring microbial genera in a conventionally-managed experimental field in the south-east of The Netherlands is remarkable.

Some nematophagous fungi such as *Arthrobotrys oligospora* have been recorded from various continents and in a wide range of habitats, including agricultural fields (Niu and Zhang, 2011). Hence, ample representation of this facultative nematophagous genus was to be expected. Essentially the same holds for *Nematoctonus leiosporus,* the most frequently isolated endoparasite among the Basidiomycota (Gray, 1983). On the other hand, the nematode antagonist *Mortierella globalpina* has been isolated and characterized only recently (DiLegge et al., 2019), and virtually nothing is known about its global distribution. From the few available reports on the endoparasite *Haptocillium balanoides* (also known as *Verticillium balanoides*) it may be concluded that the species is distributed over multiple continents (Watanabe, 2000; Farrell et al., 2006). Hence, the putative nematode antagonist community detected in the Vredepeel experimental field comprised species which presence could be anticipated supplemented with taxa that we did not foresee to be present. *Pochonia chlamydosporia,* one of the most studied nematophagous fungi, was not found in our experimental field. This is noteworthy as it has been reported as an abundant and widespread naturally occurring egg-parasite of *Meloidogyne* in organic and integrated vegetable production systems (Giné et al., 2013; Ghahremani et al., 2022).

### Soil nutritional condition determines the trophic lifestyle of most nematode antagonists

Although some antagonists have an obligate nematophagous lifestyle such as *Hirsutella vermicola* (Xiang et al., 2006) and *Drechmeria coniospora* (Lebrigand et al., 2016), most nematophagous microbes are facultative nematode antagonists. The switch from a saprophytic to a nematophagous lifestyle is triggered by the environment. Physical contact between hyphae of the facultative nematophagous fungus *Arthrobotrys oligospora* and a nematode was shown to constitute a strong trigger for trap formation (Tunlid et al., 1992), but the actual response is co-determined by N and C availability. On substrates without nitrogen, traps were formed irrespectively of the C availability, and limiting the carbon availability in the presence of nitrogen resulted in a stronger nematophagous response of *A. oligospora* (Scholler and Rubner, 1994). Although this nutritional response has mainly been characterized for *A. oligospora*, we expect that most facultative nematode antagonists will respond to the soil environment in a comparable way. We characterized the soil nutritional status of the Vredepeel experimental field (Materials and methods 2.1), and with a C/N ratio = 18 – 21, and a N content of ≈ 4,000 kg N ha^-1^ this soil had a nutritional status that is unlikely to trigger a switch from a saprophytic to a nematophagous lifestyle among facultative nematode antagonists.

### Impact of cover crops on the local nematode antagonist’ community

The growing of cover crops caused significant shifts in the community structure, diversity and abundance of putative nematode antagonists. Oilseed radish for example strongly promoted the resident *Pseudomonas* (γ-proteobacteria) community (Table 5), and whereas the oilseed radish cultivar Terranova induced an increase of *Arthrobotrys* spp. at DNA level, in combination with vetch it had the highest impact on the activity of representatives of this genus (Table 5). From our results it can be concluded that cover crops have distinct effects on microbial genera harbouring nematode antagonists, and – in addition - cover crop treatments that promote the presence of given genus, do not necessarily have a positive effect on the activity of the members of the same genus.

We observed intraspecific variation regarding stimulation of nematode antagonists in oilseed radish. Comparisons of the impacts of monocultures of the cultivars Adios, Radical, Terranova on *Arthrobotrys* spp. revealed that Terranova had the strongest stimulating effect on antagonist density, whereas activity was more promoted by the cultivars Terranova and Radical than by the cultivar Adios. This observation suggests for genotypic variation in oilseed radish with regard to the steering of nematode antagonists. Genotypic variation among oilseed radish cultivars on the soil microbiome in general was previously shown in (Cazzaniga et al., 2023a; Cazzaniga et al., 2023b).

Intraspecific variation regarding microbiome selection has been shown before for various plant species. In case of tomato, genotypic variation accounted for 10% of the variation in root microbiota (French et al., 2020; Oyserman et al., 2022). In another study, the effect of domesticated and wild barley varieties on the soil microbiome was tested, and elite varieties exerted a stronger effect on the rhizobiome than the non-domesticated genotypes (Alegria Terrazas et al., 2020). It is concluded that cover crops can be used to stimulate microbial genera of putative nematode antagonists, but they do so in a species- and even genotype-specific manner.

### Impact of *M. chitwoodi* density on the local antagonist community

Only a few putative nematode-antagonistic genera responded to differences in *M. chitwoodi* densities. Whereas *Bacillus* (Firmicutes) and *Nematoctonus* (Basidiomycota) were apparently unaffected by the densities of this plant-parasitic nematode, other fungal genera such as *Trichoderma and Arthrobotrys* responded to cover crop treatments only in plots with the highest *M. chitwoodi* densities. The genus *Haptocillium* constituted the clearest example of a genus harbouring nematode antagonists showing higher abundances in plots with elevated *M. chitwoodi* irrespective of the cover crop treatment (Table 5). This can – at least in part - be explained by its “low saprophytic ability”, that was reported for one of the detected species, *H. balanoides* (Zare and Games, 2003). No similar ecological information could be found about the second species, *H. sinense* (Table 5). Hence, the increased presence of the (semi-) obligate nematophagous fungal genus *Haptocillium* can probably best be explained by its trophic ecology. Another significant effect of *M. chitwoodi* density was observed for the fungal genus *Mortierella,* represented in the Vredepeel experimental field with three species. *M. globalpina* was recently identified as a potential biocontrol agent against *M. chitwoodi* (DiLegge et al., 2019). Virtually no ecological information is available for *M. elongatula* and *M. fatshederae.* It is concluded that *M. chitwoodi* density stimulated a few local putative nematode antagonists, but the effect of nematode density was less pronounced than the effect of cover crop treatments.

### Limitations regarding the identification of putative nematode antagonists

The diversity of putative nematode antagonists found in this study is likely an underestimation of the overall diversity of antagonists in the field, because over time more ecological functions of soil bacteria and fungi will be discovered. The two authoritative reviews on antagonists of plant-parasitic nematodes only provided a clear starting point for our investigation, showing the potential of local microbiota in the field.

Illumina NovaSeq (2x 250 bp) was used to characterize microbial amplicons. Upon sequencing ≈ 500 bp per microbial genotype, a fraction of the reads could be accurately assigned to the genus level (17% and 19% for bacteria and fungi, respectively), and ∼5% of bacteria and fungi could be classified until species level. Due to the massive output of NovaSeq sequencing (around 1 billion reads per SP flow cell), these are still considerable numbers. Hence, especially at the species level, we might have missed antagonists. As a consequence, the richness in putative nematode antagonists present in the arable field under investigation could be higher than currently reported.

## 5. Conclusion and future outlook

Antagonists of plant-parasitic nematodes have been studied for decades, and numerous examples of nematode antagonistic bacteria and fungi have been documented (Li et al., 2015; Topalovic et al., 2020). These efforts have provided important insights in the diverse and often elegant mechanisms by which microbial antagonist predate on nematodes. However, these insights could seldomly be translated into effective biocontrol measures as introduction of biocontrol agents in soil – a highly competitive environment – appeared often problematic. In this study, we focused on ways to stimulate the local nematode-antagonistic community in an arable soil. We found a strikingly large richness of putative nematode antagonists in this system. We demonstrated that bacterial and fungal genera comprising plant-parasitic nematode antagonists can be steered by cover crops, and – to a lesser extent – by plant-parasitic nematode density. Notably, none of the two variables had a generic positive or negative effect on the conglomerate of local putative nematode antagonists. A set of microbial taxa harbouring nematode antagonists were promoted in a cover crop-specific manner. We also demonstrated that some nematode antagonists were stimulated by cover crops in the presence of high *M. chitwoodi* densities only. If common arable soils indeed harbour a more diverse nematode-antagonistic community than previously anticipated, future research should focus on the identification of cues that promote the switch from a saprophytic to a nematophagous lifestyle for a relevant fraction of local nematode antagonistic. community.

## 6. Acknowledgements

The authors would like to acknowledge all the partners of the PPS Groenbemeters: LTO Nederland, Van Iperen B.V., BO Akkerbouw, Tree Centre Opheusden (TCO) and Barenbrug, DSV Zaden, Joordens Zaden and Vandinter Semo for their help in selecting and providing the cover crops seeds used in the experiment; Harry Verstegen, Marc Kroonen and the Vredepeel field team for setting up and maintaining the field experiment. We also like to thank Davide Francioli, Eline Ampt, Francesco Garassino, Lize Braat, Paula Harkes, Silvia Calderone and Thimo Peters for helping with the field samplings. We thank Frank Becker for his help with the preparation of the samples for NovaSeq sequencing. We acknowledge the Utrecht Sequencing Facility (USEQ) for providing sequencing service and data. USEQ is subsidized by the University Medical Center Utrecht and The Netherlands X-omics Initiative (NWO project 184.034.019).

