## Supplementary material for "On the diversity of nematode antagonists in an agricultural soil, and their steerability by root-knot nematode density and cover crops": Suppl Tables 1-4

### Supplementary Tables

**Suppl. Table 1 |** Table of genera harbouring putative nematode antagonists considered in this study based on two authoritative reviews (Topalovic et al (2020), Li et al (2015b)). For each genus, the species representation in the Vredepeel experimental field is presented. This includes all representatives, this table includes species that show no interaction with plant-parasitic nematodes, or species for which no relevant ecological information is available. Prevalences, the number of soil samples comprising representatives of a given genus, are presented for T0 and T1 at DNA and at RNA level.

| | **Bacterial genera** | Source | Representatives of genus found in Vredepeel experimental field | DNA-T0 Prevalence (12 samples) | RNA-T0 Prevalence (12 samples) | DNA-T1 Prevalence (129 samples) | RNA-T1 Prevalence (130 samples) |
|---|---|---|---|---|---|---|---|
| 1 | *Arthrobacter* | A | | | | | |
| 2 | *Bacillus* | A, B | *B. asahii, B. licheniformis, B. longiquaesitum, B. murimartini, B. oleronius, B.plakortidis, B. thermoamylovorans, B. smithii* | 12 | 11 | 125 | 124 |
| 3 | *Lysobacter* | A | *L. concretionis, L. dokdonensis, L. ginsengisoli, L. niastensis, L. soli* | 12 | 12 | 129 | 129 |
| 4 | *Pasteuria* | A, B | | | | | |
| 5 | *Pseudomonas* | A, B | *P. putida, P. stutzeri, P. alcaligenes, P. caeni* | 12 | 12 | 127 | 128 |
| 6 | *Rhizobium* | A | *R. arenae, R. azooxidifex, R. giardinii, R. leguminosarum, R. mongolense, R. soli, R. yantingense* | 12 | 12 | 124 | 129 |
| 7 | *Streptomyces* | A | | | | | |
| 8 | *Variovorax* | A, B | *V. paradoxus, V. soli* | 12 | 12 | 127 | 127 |

| | ***Fungal genera*** | Source | Representatives of genus found in Vredepeel experimental field | DNA-T0 Prevalence (12 samples) | RNA-T0 Prevalence (12 samples) | DNA-T1 Prevalence (130 samples) | RNA-T1 Prevalence (129 samples) |
|---|---|---|---|---|---|---|---|
| 1 | *Arthrobotrys / Orbilia* | A, B | *A. brochopaga, A. dactyloides, A. iridis, A. oligospora, A. oudemansii, A. reticulata* | 12 | 7 | 126 | 112 |
| 2 | *Dactylella* | A, B | *D. tenuis* | 1 | | 4 | 5 |
| 3 | *Dactylellina / Orbilia* (formerly *Monacrosporium*) | A, B | *M. drechsleri* | 1 | | 18 | 11 |
| 4 | *Drechslerella / Orbilia* | A, B | | | | | |
| 5 | *Orbilia* | A | *O. rectispora* | | | 3 | 2 |
| 6 | *Harposporium / Podocrella* | B | | | | | |
| 7 | *Drechmeria* | B | *D. coniospora* | 1 | | 1 | |
| 8 | *Haptocillium / Cordyceps* | A, B | *H. balanoides, H. sinense* | 5 | | 30 | 1 |
| 9 | *Hirsutella* | A, B | *H. vermicola* | 5 | | 46 | 3 |
| 10 | *Pochonia / Metacordyceps* | A, B | | | | | |
| 11 | *Purpureocillium* (formerly *Paecilomyces*) | A, B | | | | | |
| 12 | *Lecanicillium / Cordyceps* | B | | 1 | | | |
| 13 | *Trichoderma* | A, B | *T. albolutescens* | 12 | 10 | 124 | 85 |
| 14 | *Acremonium* | B | *A. biseptum, A. furcatum, A. rutilum, A. verruculosum* | 8 | | 55 | 13 |
| 15 | *Neotyphodium* | B | | | | | |
| 16 | *Fusarium* | B | *F. poae* | 12 | 6 | 130 | 106 |
| 17 | *Penicillium* | B | [No resolution beyond genus level] | 12 | 6 | 130 | 44 |
| 18 | *Nematoctonus / Hohenbuehelia* | B | *N. leiosporus* | 8 | | 34 | 6 |
| 19 | *Pleurotus* | B | | | | | |
| 20 | *Coprinus + Coprinellus* | B | *C. heptemerus* | 2 | 1 | 17 | 4 |
| 21 | *Stropharia* | B | | | | 1 | |

| | | | | | | | |
|---|---|---|---|---|---|---|---|
| 22 | *Glomus* | B | | | | | |
| 23 | *Catenaria* | A, B | | | | | |
| 24 | *Stylopage* | B | | | | | |
| 25 | *Mortierella* | A | *M. elongatula, M. fatshederae, M. globalpina* | 12 | 9 | 116 | 93 |
| 26 | *Cystopage* | B | | | | | |

9

**Suppl. Tables 2 |** PERMANOVA analyses on the bacterial and fungal communities at T0 (A), T1 (B) and pairwise comparisons at T1 (C)

**A** PERMANOVA analysis of the bacterial and fungal community composition at T0 - during pre-crops

| | Bacteria DNA | Bacteria RNA | Fungi DNA | Fungi RNA |
|---|---|---|---|---|
| block | $R^2$ = 22.4%, P=0.001, | $R^2$=17.3%, P=0.002 | , $R^2$=38.1%, P=0.001 | $R^2$ = 15.4%, P=0.032, |
| sample type | $R^2$=18.7%, P=0.001 | $R^2$=20.6%, P=0.001, | $R^2$=14.0%, P=0.001, | $R^2$=18.0%, P=0.001, |
| pre crop | ns | ns | ns | ns |
| RKN density | ns | ns | ns | ns |

**B** PERMANOVA analysis of the bacterial and fungal community composition at T1 - cover crop rhizosphere + fallow

| | Bacteria DNA | Bacteria RNA | Fungi DNA | Fungi RNA |
|---|---|---|---|---|
| block | $R^2$ = 5.9%, P=0.001 | $R^2$=4.7%, P=0.001 | $R^2$=6.1%, P=0.001, | $R^2$=3.3%, P=0.001 |
| RKN density | ns | ns | $R^2$=2.3%, P=0.005 | $R^2$=2.2%, P=0.002 |
| cover crop | **$R^2$=22.4%, P=0.001** | **$R^2$=25.5%, P=0.001,** | **$R^2$=27.1%, P=0.001,** | **$R^2$=21.0%, P=0.001,** |
| RKN density * cover crop | ns | ns | ns | ns |

**C** Pairwise comparisons of the microbiomes associated with cover crops rhizosphere and fallow at T1. P-value adjusted with Bonferroni-Holmes method. Highlighted in green significant differences

Bacteria - DNA

| | FW | OSR_T | OSR_A | OSR_R | PHA | VE | BO | PHA.OSR_T | VE.OSR_T | BO.OSR_T |
|---|---|---|---|---|---|---|---|---|---|---|
| OSR_T | 0.0016 | - | - | - | - | - | - | - | - | - |
| OSR_A | 0.0016 | 0.567 | - | - | - | - | - | - | - | - |
| OSR_R | 0.0016 | 0.3269 | 0.5184 | - | - | - | - | - | - | - |
| PHA | 0.0016 | 0.0016 | 0.0016 | 0.0016 | - | - | - | - | - | - |
| VE | 0.0138 | 0.003 | 0.0016 | 0.0056 | 0.0016 | - | - | - | - | - |
| BO | 0.0016 | 0.0016 | 0.0016 | 0.0016 | 0.0016 | 0.0016 | - | - | - | - |
| PHA.OSR_T | 0.0016 | 0.0928 | 0.1111 | 0.0854 | 0.0056 | 0.021 | 0.0016 | - | - | - |
| VE.OSR_T | 0.0016 | 0.1745 | 0.1531 | 0.1066 | 0.0016 | 0.0016 | 0.0016 | 0.0682 | - | - |
| BO.OSR_T | 0.0016 | **0.023** | 0.0263 | 0.0161 | 0.0016 | 0.0016 | 0.0016 | 0.003 | 0.033 | - |
| BO.OSR_R | 0.0016 | 0.0016 | 0.003 | 0.0016 | 0.0016 | 0.0016 | 0.0016 | 0.0016 | 0.0016 | 0.0016 |

Bacteria - RNA

| | FW | OSR_T | OSR_A | OSR_R | PHA | VE | BO | PHA.OSR_T | VE.OSR_T | BO.OSR_T |
|---|---|---|---|---|---|---|---|---|---|---|
| OSR_T | 0.0016 | - | - | - | - | - | - | - | - | - |
| OSR_A | 0.0016 | 0.345 | - | - | - | - | - | - | - | - |
| OSR_R | 0.0016 | 0.2455 | 0.1523 | - | - | - | - | - | - | - |
| PHA | 0.0016 | 0.0016 | 0.003 | 0.0016 | - | - | - | - | - | - |
| VE | 0.0016 | 0.0016 | 0.0016 | 0.0016 | 0.0016 | - | - | - | - | - |
| BO | 0.0016 | 0.0016 | 0.0016 | 0.0016 | 0.0016 | 0.0016 | - | - | - | - |
| PHA.OSR_T | 0.0016 | 0.1806 | 0.0985 | 0.0425 | 0.003 | 0.0016 | 0.0016 | - | - | - |
| VE.OSR_T | 0.0016 | 0.1122 | 0.1154 | 0.0477 | 0.0016 | 0.0016 | 0.0016 | 0.0971 | - | - |
| BO.OSR_T | 0.0016 | 0.1033 | 0.0183 | 0.0183 | 0.0016 | 0.0016 | 0.0016 | 0.0789 | 0.0183 | - |
| BO.OSR_R | 0.0016 | **0.013** | 0.0183 | **0.0016** | 0.0016 | 0.0016 | 0.0016 | 0.0016 | 0.0016 | 0.0217 |

FUNGI - DNA

| | FW | OSR_T | OSR_A | OSR_R | PHA | VE | BO | PHA.OSR_T | VE.OSR_T | BO.OSR_T |
|---|---|---|---|---|---|---|---|---|---|---|
| OSR_T | 0.0016 | - | - | - | - | - | - | - | - | - |
| OSR_A | 0.0016 | 0.5113 | - | - | - | - | - | - | - | - |
| OSR_R | 0.0016 | 0.3196 | 0.689 | - | - | - | - | - | - | - |
| PHA | 0.0016 | 0.0016 | 0.0016 | 0.0016 | - | - | - | - | - | - |
| VE | 0.0016 | 0.0016 | 0.0016 | 0.0016 | 0.0016 | - | - | - | - | - |
| BO | 0.0016 | 0.0016 | 0.0016 | 0.0016 | 0.0016 | 0.0016 | - | - | - | - |
| PHA.OSR_T | 0.0016 | **0.0358** | 0.2729 | 0.1914 | 0.0016 | 0.0016 | 0.0016 | - | - | - |
| VE.OSR_T | 0.0016 | 0.248 | 0.1897 | 0.165 | 0.0016 | 0.0043 | 0.0016 | 0.0375 | - | - |
| BO.OSR_T | 0.0016 | 0.0587 | 0.1369 | 0.0069 | 0.0016 | 0.0016 | 0.0016 | 0.0157 | 0.0861 | - |

| | | | | | | | | | | |
|---|---|---|---|---|---|---|---|---|---|---|
| BO.OSR_R | 0.0016 | **0.003** | 0.0094 | **0.0016** | 0.003 | 0.0016 | 0.0016 | 0.0016 | 0.0016 | 0.0069 |

FUNGI - RNA

| | FW | OSR_T | OSR_A | OSR_R | PHA | VE | BO | PHA.OSR_T | VE.OSR_T | BO.OSR_T |
|---|---|---|---|---|---|---|---|---|---|---|
| OSR_T | 0.0018 | - | - | - | - | - | - | - | - | - |
| OSR_A | 0.0018 | 0.809 | - | - | - | - | - | - | - | - |
| OSR_R | 0.0018 | 0.4717 | 0.4325 | - | - | - | - | - | - | - |
| PHA | 0.0018 | 0.0018 | 0.0031 | 0.0018 | - | - | - | - | - | - |
| VE | 0.0018 | 0.0018 | 0.0046 | 0.0018 | 0.0018 | - | - | - | - | - |
| BO | 0.0018 | 0.0018 | 0.0018 | 0.0018 | 0.0018 | 0.0018 | - | - | - | - |
| PHA.OSR_T | 0.0018 | 0.0859 | 0.5583 | 0.0217 | 0.0031 | 0.0031 | 0.0018 | - | - | - |
| VE.OSR_T | 0.0018 | 0.1351 | 0.1785 | 0.1162 | 0.0018 | 0.0018 | 0.0018 | 0.089 | - | - |
| BO.OSR_T | 0.0018 | 0.0985 | 0.662 | 0.0591 | 0.0018 | 0.0018 | 0.0018 | 0.1271 | 0.2849 | - |
| BO.OSR_R | 0.0018 | 0.0018 | 0.1381 | **0.0018** | 0.0059 | 0.0018 | 0.0018 | 0.1581 | 0.0031 | 0.0381 |

**Suppl. Table 3 |** PERMANOVA analyses on the community of bacterial and fungal antagonists at T0 (A), T1 (B) and pairwise comparisons at T1 (C)

**A** PERMANOVA analysis of the antagonists community composition at T0 - during pre-crops

| | Bacteria DNA | Bacteria RNA | Fungi DNA | Fungi RNA |
|---|---|---|---|---|
| block | ns | R2=15,6%, P=0.035 | R2=20.0%, P=0.005 | R2=20.5%, P=0.006 |
| sample type | R2=49%, P=0.001 | R2=25.8%, P=0.001 | R2=12.4%, P=0.001 | R2=9.7%, P=0.005 |
| pre crop | ns | ns | ns | ns |
| RKN density | ns | ns | ns | ns |

**B** PERMANOVA analysis of the antagonists community composition at T1 - in cover crop rhizosphere

| | Bacteria DNA | Bacteria RNA | Fungi DNA | Fungi RNA |
|---|---|---|---|---|
| block | R2=3.1%, P=0.003 | R2 =4.6%, P=0.001 | R2=3.7%, P=0.023 | ns |
| initial Mc levels | ns | ns | R2=2.8%, P=0.027 | ns |
| cover crop | R2=30.7%, P=0.001 | R2=19.3%, P=0.001 | R2=14.2%, P=0.001 | R2=9.5%, P=0.027 |
| RKN density * cover crop | ns | ns | ns | ns |

**C** Pairwise comparisons of the communities of antagonists associated with cover crops rhizosphere and fallow at T1. P-value adjusted with Bonferroni -Holmes method

Bacteria _DNA

| | FW | OSR_T | OSR_A | OSR_R | PHA | VE | BO | BO.OSR_R | BO.OSR_T | PHA.OSR_T |
|---|---|---|---|---|---|---|---|---|---|---|
| OSR_T | 0.002 | - | - | - | - | - | - | - | - | - |
| OSR_A | 0.002 | 0.392 | - | - | - | - | - | - | - | - |
| OSR_R | 0.002 | 0.265 | 0.376 | - | - | - | - | - | - | - |
| PHA | 0.002 | 0.002 | 0.002 | 0.002 | - | - | - | - | - | - |
| VE | 0.002 | 0.003 | 0.002 | 0.002 | 0.002 | - | - | - | - | - |
| BO | 0.002 | 0.002 | 0.002 | 0.002 | 0.002 | 0.002 | - | - | - | - |
| BO.OSR_R | 0.002 | 0.052 | 0.219 | 0.013 | 0.003 | 0.002 | 0.002 | - | - | - |
| BO.OSR_T | 0.002 | 0.397 | 0.136 | 0.204 | 0.002 | 0.002 | 0.002 | 0.006 | - | - |
| PHA.OSR_T | 0.002 | 0.370 | 0.332 | 0.906 | 0.003 | 0.003 | 0.002 | 0.009 | 0.280 | - |
| VE.OSR_T | 0.002 | 0.392 | 0.025 | 0.013 | 0.002 | 0.002 | 0.002 | 0.003 | 0.081 | 0.043 |

Bacteria - RNA

| | FW | OSR_T | OSR_A | OSR_R | PHA | VE | BO | BO.OSR_R | BO.OSR_T | PHA.OSR_T |
|---|---|---|---|---|---|---|---|---|---|---|
| OSR_T | 0.003 | - | - | - | - | - | - | - | - | - |
| OSR_A | 0.003 | 0.721 | - | - | - | - | - | - | - | - |
| OSR_R | 0.003 | 0.165 | 0.370 | - | - | - | - | - | - | - |
| PHA | 0.003 | 0.030 | 0.121 | 0.003 | - | - | - | - | - | - |
| VE | 0.003 | 0.003 | 0.003 | 0.003 | 0.003 | - | - | - | - | - |
| BO | 0.003 | 0.033 | 0.013 | 0.005 | 0.013 | 0.003 | - | - | - | - |

| | | | | | | | | | | |
|---|---|---|---|---|---|---|---|---|---|---|
| BO.OSR_R | 0.003 | 0.357 | 0.185 | 0.003 | 0.195 | 0.003 | 0.020 | - | - | - |
| BO.OSR_T | 0.003 | 0.467 | 0.152 | 0.299 | 0.036 | 0.003 | 0.343 | 0.042 | - | - |
| PHA.OSR_T | 0.005 | 0.259 | 0.661 | 0.357 | 0.399 | 0.003 | 0.078 | 0.034 | 0.394 | - |
| VE.OSR_T | 0.003 | 0.440 | 0.357 | 0.152 | 0.015 | 0.003 | 0.005 | 0.024 | 0.259 | 0.399 |

Fungi - DNA

| | FW | OSR_T | OSR_A | OSR_R | PHA | VE | BO | BO.OSR_R | BO.OSR_T | PHA.OSR_T |
|---|---|---|---|---|---|---|---|---|---|---|
| OSR_T | 0.037 | - | - | - | - | - | - | - | - | - |
| OSR_A | 0.028 | 0.401 | - | - | - | - | - | - | - | - |
| OSR_R | 0.087 | 0.168 | 0.401 | - | - | - | - | - | - | - |
| PHA | 0.067 | 0.179 | 0.401 | 0.168 | - | - | - | - | - | - |
| VE | 0.168 | 0.345 | 0.399 | 0.179 | 0.321 | - | - | - | - | - |
| BO | 0.028 | 0.110 | 0.241 | 0.168 | 0.391 | 0.082 | - | - | - | - |
| BO.OSR_R | 0.033 | 0.231 | 0.932 | 0.401 | 0.389 | 0.287 | 0.343 | - | - | - |
| BO.OSR_T | 0.033 | 0.062 | 0.343 | 0.168 | 0.343 | 0.039 | 0.766 | 0.401 | - | - |
| PHA.OSR_T | 0.168 | 0.406 | 0.401 | 0.343 | 0.458 | 0.814 | 0.102 | 0.310 | 0.085 | - |
| VE.OSR_T | 0.033 | 0.168 | 0.440 | 0.310 | 0.736 | 0.168 | 0.401 | 0.401 | 0.453 | 0.310 |

Fungi - RNA

| | FW | OSR_T | OSR_A | OSR_R | PHA | VE | BO | BO.OSR_R | BO.OSR_T | PHA.OSR_T |
|---|---|---|---|---|---|---|---|---|---|---|
| OSR_T | 0.780 | - | - | - | - | - | - | - | - | - |
| OSR_A | 0.700 | 0.730 | - | - | - | - | - | - | - | - |
| OSR_R | 0.560 | 0.460 | 0.520 | - | - | - | - | - | - | - |
| PHA | 0.460 | 0.520 | 0.900 | 0.670 | - | - | - | - | - | - |
| VE | 0.640 | 0.520 | 0.770 | 0.320 | 0.460 | - | - | - | - | - |
| BO | 0.210 | 0.210 | 0.720 | 0.210 | 0.500 | 0.520 | - | - | - | - |
| BO.OSR_R | 0.180 | 0.210 | 0.600 | 0.180 | 0.460 | 0.460 | 0.520 | - | - | - |
| BO.OSR_T | 0.460 | 0.400 | 0.940 | 0.730 | 0.730 | 0.520 | 0.720 | 0.600 | - | - |
| PHA.OSR_T | 0.600 | 0.800 | 0.980 | 0.700 | 0.980 | 0.860 | 0.720 | 0.730 | 0.940 | - |
| VE.OSR_T | 0.180 | 0.210 | 0.770 | 0.460 | 0.900 | 0.460 | 0.460 | 0.210 | 0.560 | 0.770 |

**Suppl. Table 4 |** Results of the generalised negative binomial mixed models on the counts of putative nematode antagonists. Post-hoc tests were conducted with emmeans and BH procedure for multiple comparisons

Fungi

Acremonium

DNA

| cc | response | SE | .group |
|---|---|---|---|
| OSR_T | 45.8 | 15.1 | a |
| OSR_A | 65.5 | 23.1 | a |
| OSR_R | 221 | 133.6 | ab |
| PHA | 44.7 | 17.2 | a |
| VE | 59.4 | 25.7 | a |
| BO | 53.6 | 14.6 | a |
| BO.OSR_R | 85.1 | 27.7 | a |
| BO.OSR_T | 62.3 | 26.8 | a |
| PHA.OSR_T | 69 | 41 | a |
| VE.OSR_T | 676.1 | 257.6 | b |

RNA

no model, prevalence too low

Arthrobotrys

DNA

RKN level Low

| cover crop | response | SE | .group |
|---|---|---|---|
| OSR_T | 3133.8 | 1712.8 | d |
| OSR_A | 268.1 | 146.9 | abc |
| OSR_R | 638 | 349 | bcd |
| PHA | 87.6 | 48 | a |
| VE | 458.3 | 250.9 | abc |
| BO | 1316.8 | 719.9 | cd |
| BO.OSR_R | 275.5 | 151 | abc |
| BO.OSR_T | 163.2 | 89.5 | ab |
| PHA.OSR_T | 145.9 | 98.1 | ab |
| VE.OSR_T | 170.6 | 93.6 | ab |

RKN level Medium:

| cover crop | response | SE | .group |
|---|---|---|---|
| OSR_T | 312.7 | 121.2 | bc |
| OSR_A | 496.1 | 192 | cd |
| OSR_R | 212.4 | 82.6 | bc |

RNA

RKN level Low

| cover crop | response | SE | .group |
|---|---|---|---|
| OSR_T | 2582.7 | 2386.6 | b |
| OSR_A | 1493.9 | 1380.8 | b |
| OSR_R | 2682.6 | 2478.9 | b |
| PHA | 14.6 | 13.6 | a |
| VE | 309.6 | 286.2 | ab |
| BO | 407 | 376.5 | ab |
| BO.OSR_R | 963.1 | 890.2 | b |
| BO.OSR_T | 435.6 | 402.6 | ab |
| PHA.OSR_T | 749.9 | 848.9 | ab |
| VE.OSR_T | 253.1 | 234 | ab |

RKN level Medium:

| cover crop | response | SE | .group |
|---|---|---|---|
| OSR_T | 656.3 | 429.3 | ab |
| OSR_A | 1011.1 | 660.8 | ab |
| OSR_R | 102.9 | 67.4 | a |

| | | | | | | | | | |
|---|---|---|---|---|---|---|---|---|---|
| | PHA | 49.7 | 21.1 | a | | PHA | 427.6 | 279.6 | ab |
| | VE | 370.3 | 143.4 | bc | | VE | 734.1 | 479.8 | ab |
| | BO | 1530.8 | 591.8 | d | | BO | 1510.1 | 986.8 | ab |
| | BO.OSR_R | 267.6 | 103.7 | bc | | BO.OSR_R | 536.4 | 384 | ab |
| | BO.OSR_T | 161.7 | 62.7 | abc | | BO.OSR_T | 818 | 534.8 | ab |
| | PHA.OSR_T | 110.2 | 47.8 | ab | | PHA.OSR_T | 401.7 | 287.7 | ab |
| | VE.OSR_T | 471.1 | 182.2 | cd | | VE.OSR_T | 2212 | 1445.4 | b |
| RKN level | High: | | | | RKN level | High: | | | |
| | cover crop | response | SE | .group | | cover crop | response | SE | .group |
| | OSR_T | 3516.6 | 1922.1 | c | | OSR_T | 5820.3 | 5378.2 | b |
| | OSR_A | 220 | 120.6 | b | | OSR_A | 65 | 60.3 | a |
| | OSR_R | 416.3 | 280.1 | b | | OSR_R | 983.9 | 909.4 | ab |
| | PHA | 28.7 | 15.9 | a | | PHA | 46.1 | 42.8 | a |
| | VE | 571.2 | 312.4 | bc | | VE | 728.7 | 673.5 | ab |
| | BO | 380.4 | 208.2 | b | | BO | 1192.3 | 1101.9 | ab |
| | BO.OSR_R | 207.2 | 113.6 | b | | BO.OSR_R | 555.7 | 513.8 | ab |
| | BO.OSR_T | 847.7 | 463.8 | bc | | BO.OSR_T | 2608.1 | 2410.1 | b |
| | PHA.OSR_T | 347.1 | 190.1 | b | | PHA.OSR_T | 559.1 | 527.5 | ab |
| | VE.OSR_T | 1088.2 | 595 | bc | | VE.OSR_T | 12101.3 | 11182 | b |

**Fusarium**

| DNA | | | | | RNA | | | | |
|---|---|---|---|---|---|---|---|---|---|
| RKN level: | Low | | | | RKN level: | Low | | | |
| | cover crop | response | SE | .group | | cover crop | response | SE | .group |
| | OSR_T | 1584 | 525 | a | | OSR_T | 865 | 318 | a |
| | OSR_A | 1485 | 494 | a | | OSR_A | 637 | 285 | a |
| | OSR_R | 2199 | 731 | a | | OSR_R | 985 | 363 | a |
| | PHA | 2360 | 789 | a | | PHA | 1093 | 402 | a |
| | VE | 3828 | 1269 | a | | VE | 2436 | 896 | a |
| | BO | 3048 | 1017 | a | | BO | 1929 | 864 | a |
| | BO.OSR_R | 2090 | 699 | a | | BO.OSR_R | 1340 | 495 | a |
| | BO.OSR_T | 2752 | 913 | a | | BO.OSR_T | 1622 | 596 | a |
| | PHA.OSR_T | 4406 | 1783 | a | | PHA.OSR_T | 3903 | 1811 | a |
| | VE.OSR_T | 5849 | 1940 | a | | VE.OSR_T | 1467 | 540 | a |
| RKN: | Medium | | | | RKN: | Medium | | | |
| | cover crop | response | SE | .group | | cover crop | response | SE | .group |
| | OSR_T | 1202 | 300 | ab | | OSR_T | 1457 | 539 | a |
| | OSR_A | 2105 | 527 | abcd | | OSR_A | 1244 | 376 | a |
| | OSR_R | 2858 | 713 | cd | | OSR_R | 1491 | 437 | a |
| | PHA | 1021 | 254 | a | | PHA | 1170 | 436 | a |
| | VE | 3308 | 823 | d | | VE | 1171 | 432 | a |
| | BO | 3931 | 973 | d | | BO | 3044 | 819 | a |
| | BO.OSR_R | 2214 | 548 | abcd | | BO.OSR_R | 1844 | 540 | a |
| | BO.OSR_T | 2277 | 562 | bcd | | BO.OSR_T | 1554 | 419 | a |
| | PHA.OSR_T | 1416 | 378 | abc | | PHA.OSR_T | 1009 | 299 | a |
| | VE.OSR_T | 2350 | 583 | bcd | | VE.OSR_T | 1000 | 298 | a |
| RKN: | High | | | | RKN: | High | | | |
| | cover crop | response | SE | .group | | cover crop | response | SE | .group |
| | OSR_T | 1541 | 514 | bc | | OSR_T | 1580 | 585 | c |
| | OSR_A | 2956 | 984 | c | | OSR_A | 2066 | 922 | c |
| | OSR_R | 2019 | 674 | bc | | OSR_R | 2069 | 766 | c |
| | PHA | 345 | 115 | a | | PHA | 375 | 168 | ab |
| | VE | 2848 | 953 | c | | VE | 985 | 364 | bc |
| | BO | 3651 | 1226 | c | | BO | 1312 | 597 | bc |
| | BO.OSR_R | 868 | 289 | ab | | BO.OSR_R | 1864 | 833 | c |
| | BO.OSR_T | 2154 | 719 | bc | | BO.OSR_T | 897 | 401 | abc |
| | PHA.OSR_T | 1728 | 575 | bc | | PHA.OSR_T | 218 | 95 | a |
| | VE.OSR_T | 2007 | 667 | bc | | VE.OSR_T | 916 | 338 | bc |

**Haptocillium**

| DNA | | | | | RNA |
|---|---|---|---|---|---|
| | RKN level | response | SE | .group | no model, prevalence too low |
| | Low | 9.89 | 6.56 | a | |
| | Medium | 25.4 | 7.12 | a | |
| | High | 132.05 | 33.33 | b | |

**Hirsutella**

| DNA | RNA |
|---|---|
| ns | no model, prevalence too low |

Mortierella

DNA

ns

RNA

| | response | SE | .group |
|---|---|---|---|
| Low | 104 | 19.9 | a |
| Medium | 167 | 24.1 | b |
| High | 213 | 45.3 | b |

Nematoctonus

DNA

| cover crop | response | SE | .group |
|---|---|---|---|
| OSR_T | 9.52 | 2.85 | a |
| OSR_A | 70.74 | 24.83 | c |
| OSR_R | 35.58 | 12.75 | bc |
| PHA | 27.15 | 9.64 | abc |
| VE | 11.47 | 4.63 | ab |
| BO | 8.49 | 2.73 | a |
| BO.OSR_R | 18.64 | 5.8 | ab |
| BO.OSR_T | 15.38 | 4.72 | ab |
| PHA.OSR_T | 18.7 | 4.08 | ab |
| VE.OSR_T | 17.39 | 4.28 | ab |

RNA

no model, prevalence too low

Penicillium

DNA

| cover crop | response | SE | df |
|---|---|---|---|
| OSR_T | 635 | 132.1 | bc |
| OSR_A | 520 | 108.3 | abc |
| OSR_R | 368 | 77.3 | abc |
| PHA | 578 | 124.7 | abc |
| VE | 1503 | 313.4 | d |
| BO | 525 | 110.1 | abc |
| BO.OSR_R | 302 | 63.2 | a |
| BO.OSR_T | 347 | 72.2 | ab |
| PHA.OSR_T | 752 | 174.2 | cd |
| VE.OSR_T | 640 | 133.3 | bc |

RNA

ns

Trichoderma

DNA

| cover crop | response | SE | .group |
|---|---|---|---|
| OSR_T | 388 | 139.6 | ab |
| OSR_A | 282 | 104.7 | ab |
| OSR_R | 363 | 137.9 | ab |
| PHA | 338 | 129.3 | ab |
| VE | 666 | 239.1 | b |
| BO | 353 | 126.8 | ab |
| BO.OSR_R | 195 | 71.3 | a |
| BO.OSR_T | 214 | 77.4 | a |
| PHA.OSR_T | 245 | 93.1 | ab |
| VE.OSR_T | 373 | 134.9 | ab |

RNA

RKN: Low

| cover crop | response | SE | .group |
|---|---|---|---|
| OSR_T | 266.2 | 166.3078 | a |
| OSR_A | 61.5 | 52.6739 | a |
| OSR_R | 272.4 | 166.2404 | a |
| PHA | 437.5 | 271.2154 | a |
| VE | 153.7 | 79.2173 | a |
| BO | 111.5 | 65.3167 | a |
| BO.OSR_R | 262.4 | 133.3852 | a |
| BO.OSR_T | 424.9 | 262.0196 | a |
| PHA.OSR_T | 515.2 | 316.467 | a |
| VE.OSR_T | 242.4 | 125.9103 | a |

RKN: Medium

| cover crop | response | SE | .group |
|---|---|---|---|
| OSR_T | 1496.3 | 615.684 | b |
| OSR_A | 351 | 147.3155 | ab |
| OSR_R | 798.6 | 484.7013 | ab |
| PHA | 309.2 | 164.8237 | ab |
| VE | 536.6 | 243.6023 | ab |
| BO | 414.1 | 191.7833 | ab |
| BO.OSR_R | 106.6 | 66.3555 | a |
| BO.OSR_T | 489.6 | 220.7116 | ab |
| PHA.OSR_T | 185.3 | 94.6004 | a |
| VE.OSR_T | 470.2 | 214.5878 | ab |

RKN: High

| cover crop | response | SE | .group |
|---|---|---|---|
| OSR_T | 524.6 | 267.8484 | ab |
| OSR_A | 1214.9 | 623.7241 | b |
| OSR_R | 591.4 | 497.8063 | ab |
| PHA | 87 | 54.0437 | a |
| VE | 306.2 | 159.621 | ab |
| BO | 137.4 | 84.0664 | ab |

| | | | |
|---|---|---|---|
| BO.OSR_R | 0 | 0.0002 | ab |
| BO.OSR_T | 187.3 | 163.096 | ab |
| PHA.OSR_T | 328.8 | 194.742 | ab |
| VE.OSR_T | 194.7 | 118.9346 | ab |

#### Bacteria

##### Bacillus

DNA

RNA

RKN level: All

| cover crop | response | SE | .group |
|---|---|---|---|
| OSR_T | 436 | 81.2 | cd |
| OSR_A | 268 | 50.6 | abc |
| OSR_R | 350 | 65.7 | bcd |
| PHA | 186 | 39.6 | ab |
| VE | 326 | 63.1 | abcd |
| BO | 205 | 38.4 | ab |
| BO.OSR_R | 171 | 35.7 | a |
| BO.OSR_T | 292 | 54.8 | abcd |
| PHA.OSR_T | 295 | 59.7 | abcd |
| VE.OSR_T | 513 | 97.6 | d |

##### Lysobacter

| DNA | | | | RNA | | | |
|---|---|---|---|---|---|---|---|
| RKN level: Low | | | | | | | |
| cover crop | response | SE | .group | | response | SE | .group |
| OSR_T | 402 | 76.6 | a | OSR_T | 498 | 61.4 | a |
| OSR_A | 523 | 99.5 | ab | OSR_A | 472 | 58.6 | a |
| OSR_R | 494 | 93.8 | ab | OSR_R | 563 | 69.2 | a |
| PHA | 596 | 113.3 | ab | PHA | 511 | 68.7 | a |
| VE | 532 | 101.2 | ab | VE | 1783 | 224.8 | b |
| BO | 454 | 86.3 | a | BO | 553 | 68.4 | a |
| BO.OSR_R | 511 | 97.2 | ab | BO.OSR_R | 466 | 57.2 | a |
| BO.OSR_T | 608 | 115.4 | ab | BO.OSR_T | 550 | 70.5 | a |
| PHA.OSR_T | 1097 | 254.1 | b | PHA.OSR_T | 543 | 72.7 | a |
| VE.OSR_T | 399 | 75.9 | a | VE.OSR_T | 522 | 64.7 | a |

RKN level: Medium

| cover crop | response | SE | .group |
|---|---|---|---|
| OSR_T | 645 | 86.5 | a |
| OSR_A | 501 | 67.4 | a |
| OSR_R | 586 | 86.2 | a |
| PHA | 416 | 56.1 | a |
| VE | 566 | 76 | a |
| BO | 596 | 80 | a |
| BO.OSR_R | 556 | 74.9 | a |
| BO.OSR_T | 600 | 80.5 | a |
| PHA.OSR_T | 506 | 83.6 | a |
| VE.OSR_T | 536 | 78.8 | a |

RKN level: High

| cover crop | response | SE | .group |
|---|---|---|---|
| OSR_T | 389 | 74.2 | ab |
| OSR_A | 481 | 91.5 | abc |
| OSR_R | 669 | 126.9 | bc |
| PHA | 463 | 88 | abc |
| VE | 702 | 133 | bc |
| BO | 517 | 98.2 | abc |
| BO.OSR_R | 584 | 111 | bc |
| BO.OSR_T | 820 | 155.3 | c |
| PHA.OSR_T | 270 | 52 | a |
| VE.OSR_T | 376 | 71.8 | ab |
| | 820 | 155.3 | c |

##### Pseudomonas

| DNA | | | | RNA | | | |
|---|---|---|---|---|---|---|---|
| RKN level: Low | | | | RKN level: Low | | | |
| cover crop | response | SE | .group | cover crop | response | SE | .group |
| OSR_T | 6759 | 2381 | a | OSR_T | 439 | 69.1 | a |
| OSR_A | 4090 | 1441 | a | OSR_A | 853 | 133.5 | bc |
| OSR_R | 1934 | 682 | a | OSR_R | 586 | 91.4 | ab |
| PHA | 4282 | 1509 | a | PHA | 737 | 114.7 | abc |
| VE | 2899 | 1021 | a | VE | 1242 | 192.8 | c |

| | | | |
|---|---|---|---|
| BO | 2287 | 806 | a |
| BO.OSR_R | 3986 | 1404 | a |
| BO.OSR_T | 3290 | 1159 | a |
| PHA.OSR_T | 1186 | 512 | a |
| VE.OSR_T | 5863 | 2065 | a |

RKN level: Medium

| cover crop | response | SE | .group |
|---|---|---|---|
| OSR_T | 2345 | 584 | b |
| OSR_A | 4262 | 1062 | bcd |
| OSR_R | 5888 | 1467 | cd |
| PHA | 2953 | 736 | bc |
| VE | 450 | 112 | a |
| BO | 2623 | 654 | bc |
| BO.OSR_R | 8698 | 2167 | d |
| BO.OSR_T | 2627 | 655 | bc |
| PHA.OSR_T | 2070 | 632 | b |
| VE.OSR_T | 3477 | 866 | bc |

RKN level: High

| cover crop | response | SE | .group |
|---|---|---|---|
| OSR_T | 5797 | 2042 | bcde |
| OSR_A | 16687 | 5878 | e |
| OSR_R | 2247 | 792 | bc |
| PHA | 5090 | 1793 | bcd |
| VE | 454 | 160 | a |
| BO | 1868 | 658 | b |
| BO.OSR_R | 3142 | 1107 | bc |
| BO.OSR_T | 6390 | 2251 | cde |
| PHA.OSR_T | 6643 | 2340 | cde |
| VE.OSR_T | 10194 | 3591 | de |

| | | | |
|---|---|---|---|
| BO | 842 | 130.7 | bc |
| BO.OSR_R | 657 | 102.7 | ab |
| BO.OSR_T | 690 | 107.7 | ab |
| PHA.OSR_T | 602 | 114.9 | ab |
| VE.OSR_T | 713 | 111.3 | abc |

RKN level: Medium

| cover crop | response | SE | .group |
|---|---|---|---|
| OSR_T | 846 | 93.3 | bcd |
| OSR_A | 613 | 67.9 | ab |
| OSR_R | 614 | 67.6 | ab |
| PHA | 802 | 88.5 | bc |
| VE | 1186 | 130.5 | d |
| BO | 769 | 84.6 | bc |
| BO.OSR_R | 490 | 54.1 | a |
| BO.OSR_T | 747 | 82.3 | bc |
| PHA.OSR_T | 990 | 119.5 | cd |
| VE.OSR_T | 720 | 79.3 | bc |

RKN level: High

| cover crop | response | SE | .group |
|---|---|---|---|
| OSR_T | 745 | 116.1 | bc |
| OSR_A | 273 | 43.6 | a |
| OSR_R | 794 | 123.6 | bc |
| PHA | 713 | 136 | bc |
| VE | 1160 | 180.3 | c |
| BO | 804 | 125.1 | bc |
| BO.OSR_R | 733 | 114.4 | bc |
| BO.OSR_T | 694 | 108 | bc |
| PHA.OSR_T | 816 | 127.1 | bc |
| VE.OSR_T | 512 | 80 | b |

Rhizobium

DNA

| cover crop | response | SE | .group |
|---|---|---|---|
| OSR_T | 2297 | 246 | ab |
| OSR_A | 2527 | 267 | ab |
| OSR_R | 2452 | 262 | ab |
| PHA | 2171 | 230 | a |
| VE | 2437 | 258 | ab |
| BO | 2923 | 311 | ab |
| BO.OSR_R | 2602 | 275 | ab |
| BO.OSR_T | 3378 | 359 | b |
| PHA.OSR_T | 2377 | 281 | ab |
| VE.OSR_T | 3429 | 363 | b |

| RKN level | | | |
|---|---|---|---|
| Low | 2812 | 245 | b |
| Medium | 2309 | 164 | a |
| High | 2925 | 254 | b |

RNA

RKN level: Low

| cover crop | response | SE | .group |
|---|---|---|---|
| OSR_T | 2050 | 605 | a |
| OSR_A | 1493 | 441 | a |
| OSR_R | 1349 | 399 | a |
| PHA | 866 | 256 | a |
| VE | 904 | 267 | a |
| BO | 908 | 268 | a |
| BO.OSR_R | 1570 | 464 | a |
| BO.OSR_T | 1081 | 319 | a |
| PHA.OSR_T | 1323 | 479 | a |
| VE.OSR_T | 2842 | 839 | a |

RKN level: Medium

| cover crop | response | SE | .group |
|---|---|---|---|
| OSR_T | 1174 | 245 | a |
| OSR_A | 912 | 191 | a |
| OSR_R | 2898 | 605 | b |
| PHA | 1000 | 209 | a |
| VE | 879 | 184 | a |
| BO | 870 | 182 | a |
| BO.OSR_R | 1260 | 263 | a |
| BO.OSR_T | 1127 | 236 | a |
| PHA.OSR_T | 1027 | 235 | a |
| VE.OSR_T | 1053 | 220 | a |

RKN level: High

| cover crop | response | SE | .group |
|---|---|---|---|
| OSR_T | 1297 | 383 | a |
| OSR_A | 6418 | 1894 | c |
| OSR_R | 867 | 256 | a |
| PHA | 1647 | 596 | ab |
| VE | 896 | 265 | a |
| BO | 1427 | 422 | a |
| BO.OSR_R | 1711 | 505 | ab |
| BO.OSR_T | 1153 | 341 | a |
| PHA.OSR_T | 4327 | 1277 | bc |
| VE.OSR_T | 4424 | 1305 | bc |

Variovorax

DNA

| cover crop | response | SE | .group |
|---|---|---|---|
| OSR_T | 1074 | 106.7 | abcd |
| OSR_A | 997 | 99.7 | abcd |
| OSR_R | 878 | 87.6 | abc |
| PHA | 1147 | 114.2 | cd |
| VE | 790 | 78.8 | a |
| BO | 1009 | 100.1 | abcd |
| BO.OSR_R | 1267 | 126.5 | d |
| BO.OSR_T | 1337 | 132.8 | d |
| PHA.OSR_T | 791 | 88 | ab |
| VE.OSR_T | 1083 | 107.7 | bcd |

| RKN level | response | SE | .group |
|---|---|---|---|
| Low | 1127 | 82.1 | b |
| Medium | 906 | 55.1 | a |
| High | 1085 | 78.1 | b |

RNA

| cover crop | response | SE | .group |
|---|---|---|---|
| OSR_T | 281 | 37.8 | abc |
| OSR_A | 211 | 29.8 | a |
| OSR_R | 296 | 41.3 | abc |
| PHA | 438 | 58.9 | bc |
| VE | 455 | 60.9 | c |
| BO | 343 | 46.1 | abc |
| BO.OSR_R | 325 | 43.3 | abc |
| BO.OSR_T | 309 | 41.2 | abc |
| PHA.OSR_T | 438 | 63.9 | bc |
| VE.OSR_T | 261 | 35.1 | ab |
