## Supplementary material for "On the diversity of nematode antagonists in an agricultural soil, and their steerability by root-knot nematode density and cover crops": Suppl Figures 1-6

### Supplementary Figures

**Suppl. Figure S1 |** A. Lay out of the field experiment at Vredepeel. ‘bo’ = black oat, ‘ry’ = rye, ‘ar’ = annual ryegrass, ‘pr’ = perennial ryegrass pre-crops. Perpendicular to the longitudinal direction of the pre-crop strips 11 plots (each 6x3 m) were defined, and ten cover crop treatments were grown in them. VE= vetch, BO= black oat, PHA= phacelia, T= oilseed radish cultivar Terranova, R= oilseed radish cultivar Radical, A= oilseed radish cultivar Adios, BO-OSR-T= black oat – oilseed radish T, PHA-OSR-T= phacelia – oilseed radish T, VE-OSR-T= vetch - oilseed radish T, BO-OSR-R= black oat – oilseed radish R. FW = fallow, unplanted control. B) *M. chitwoodi* densities per block.

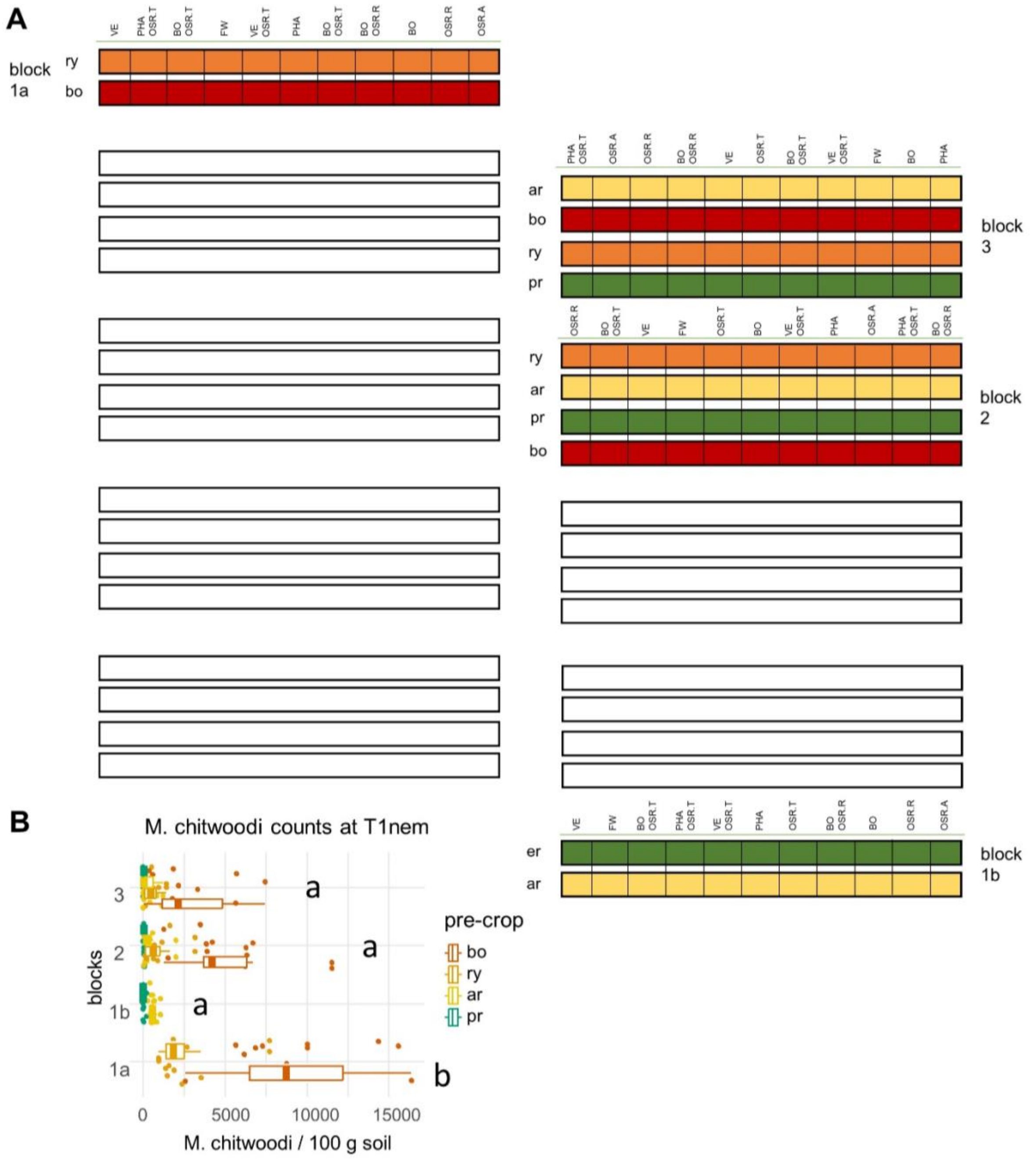

**Suppl. Figure S2 |** Principal Coordinate Analysis (PCoA) of CSS normalised ASV assigned to genera of putative nematode antagonists at T0. Dissimilarity matrix built on Bray-Curtis metric and plotted separating ASVs based on pre-crops (colours) and sample type (shapes). PERMANOVA indicated a non-significant effect of pre-crops, but a significant difference between rhizosphere ad bulk soil (Suppl. Table S2 A). The latter clearly separates samples along the principal PCoA axis.

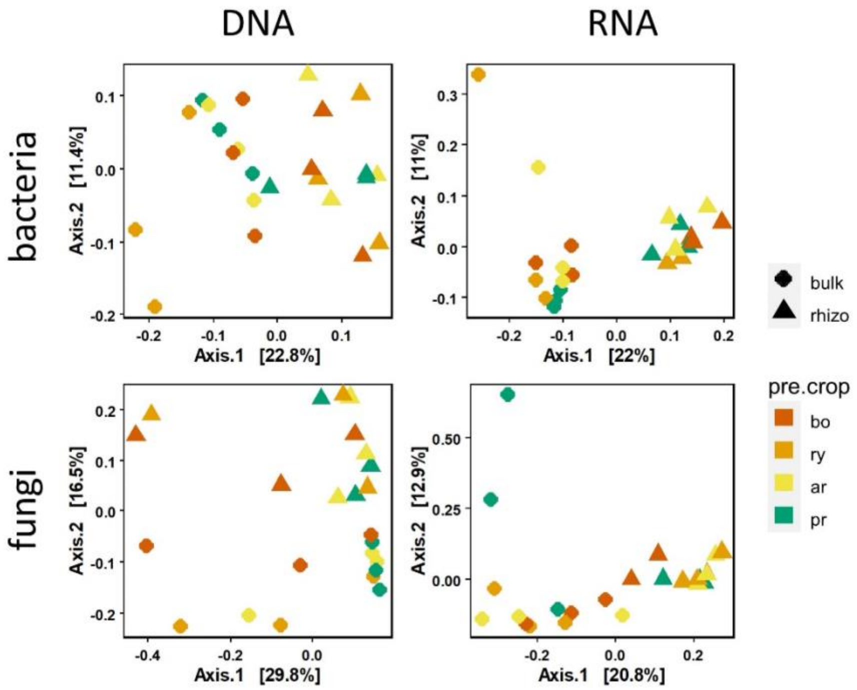

**Suppl. Figure S3 |** Diversity of putative nematode antagonists in pre-crops at T0, tested per RKN density. A and C: Principal Coordinate Analysis (PCoA) of CSS normalised ASVs assigned to bacterial and fungal putative nematode antagonists. Dissimilarity matrix built on Bray-Curtis metric and plotted separating ASVs based on RKN levels (colours). PERMANOVA indicated a non-significant effect of pre-crops. B and D: diversity (Shannon) and richness (Observed) of antagonists calculated on rarefied bacterial and fungal datasets. Differences among diversity indices per RKN level calculated with Kruskal-Wallis test and Holm adjustment for multiple testing.

Suppl. Figure 3

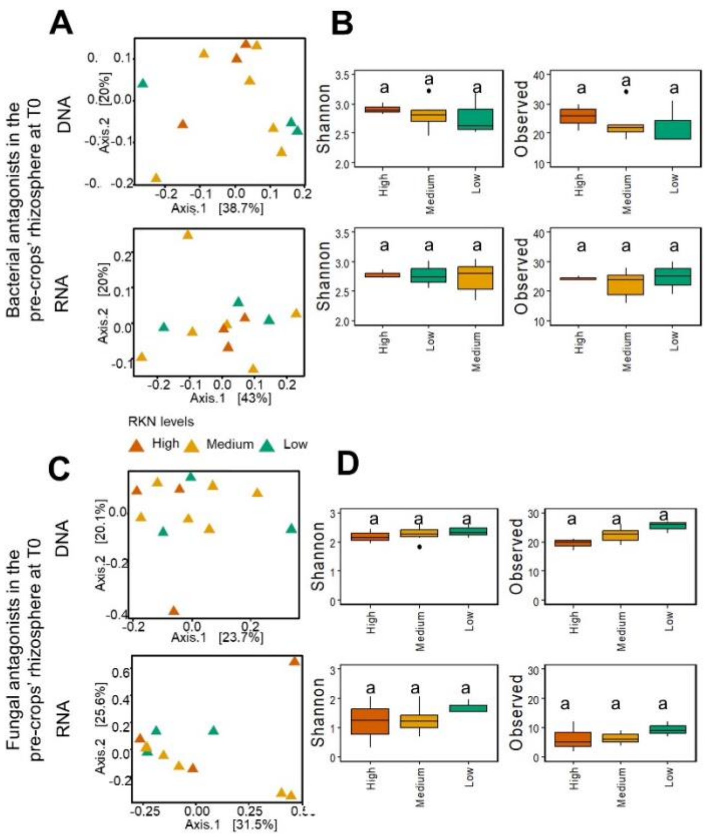

**Suppl. Figure S4 |** Principal Coordinate Analysis (PCoA) of CSS normalised ASV assigned to putative nematode-antagonistic genera at T1. Dissimilarity matrix built on Bray-Curtis metric and plotted separating ASVs based on cover crops (colours) and monocultures or mixtures (shapes). PERMANOVA indicated a significant effect of cover crops on the DNA and RNA community of bacterial and fungal putative antagonists, and a small, but significant effect of RKN levels on the fungal fraction (Suppl. Table S3 B).

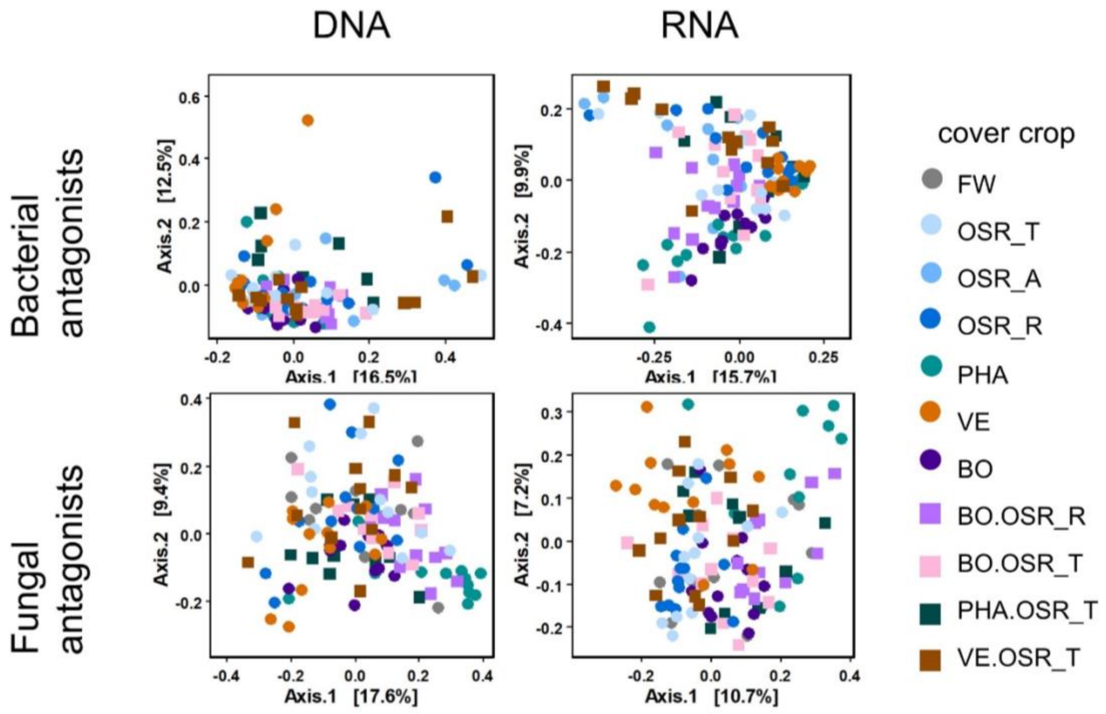

**Suppl. Figure S5 |** Diversity (Shannon) and richness (Observed) of putative nematode antagonists in the rhizosphere of cover crops and fallow at T1. A) putative bacterial antagonists at the DNA level, B) putative bacterial antagonists at the RNA level, C) putative fungal antagonists at the DNA level. No significant effect of cover crops was found on the putative fungal antagonists at the RNA level. Differences among diversity indices per cover crop treatment calculated with Kruskal-Wallis test and Holm adjustment for multiple testing.

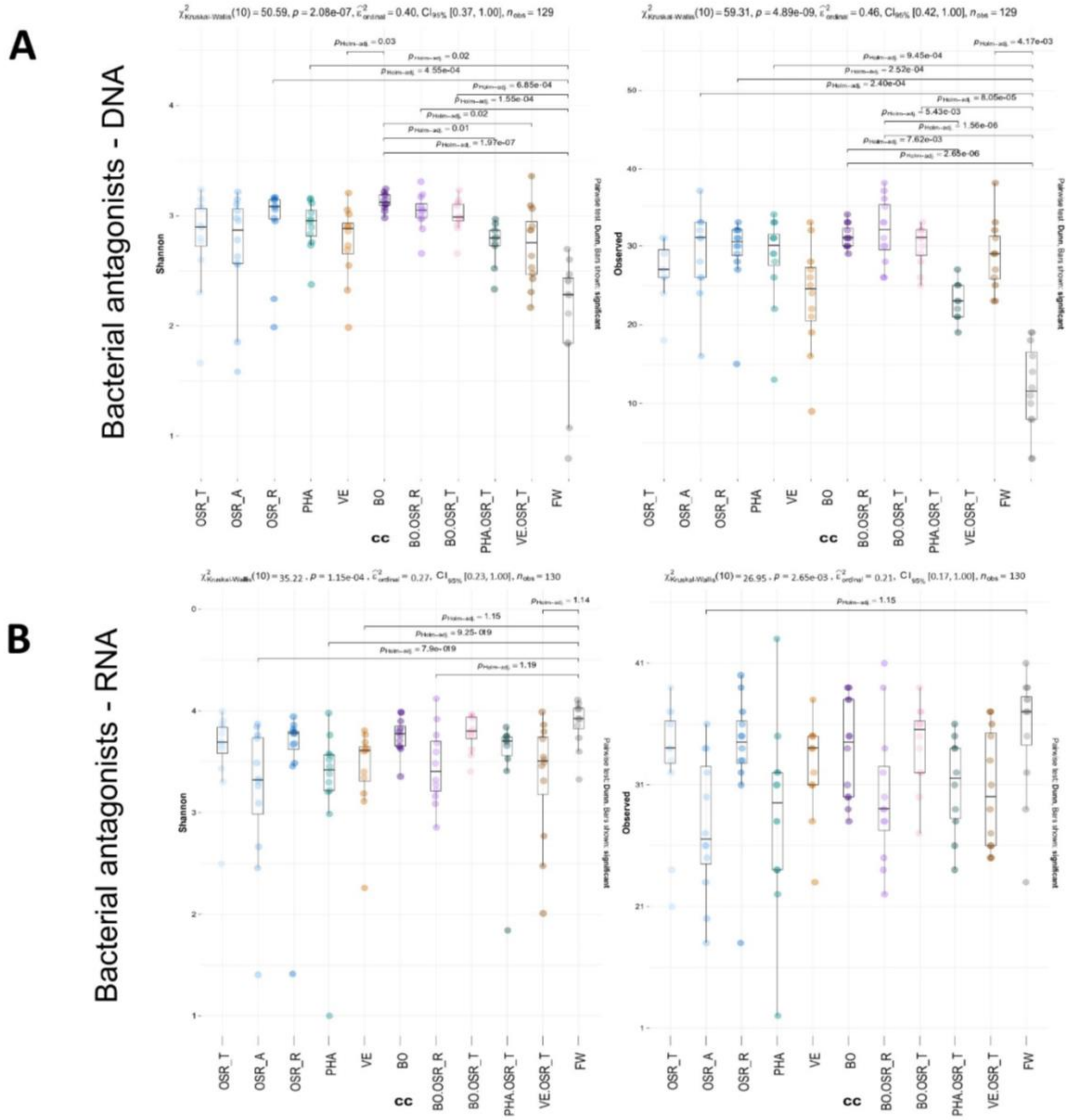

**C**

Fungal antagonists - DNA

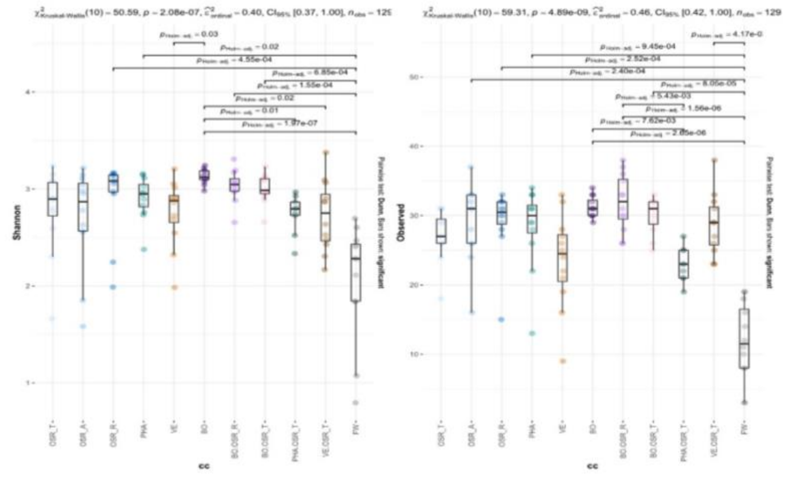

**Suppl. Figure S6 |** Relative abundance of putative nematode antagonists in cover crops rhizosphere at T1.

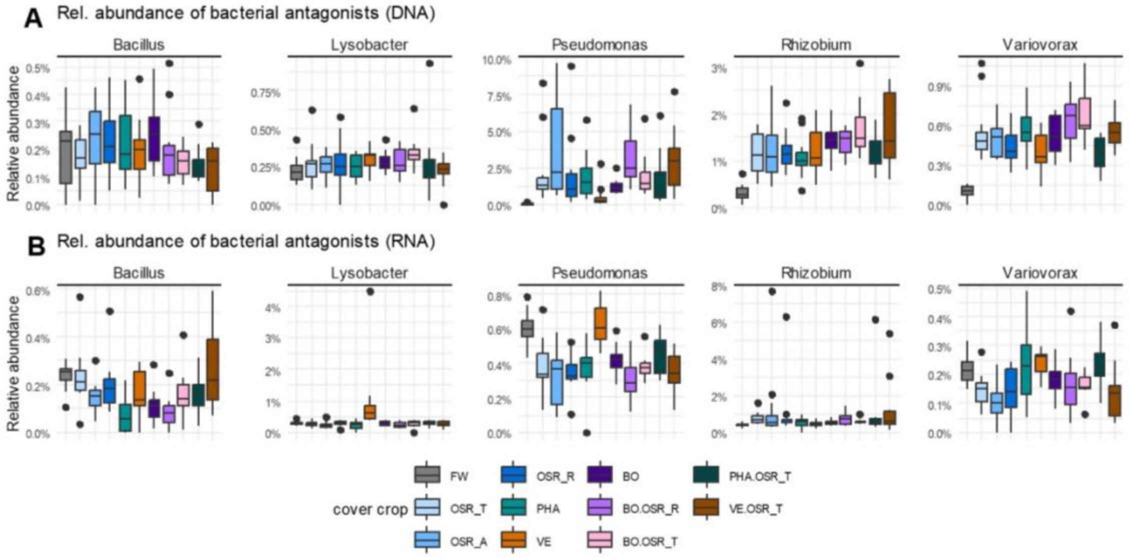


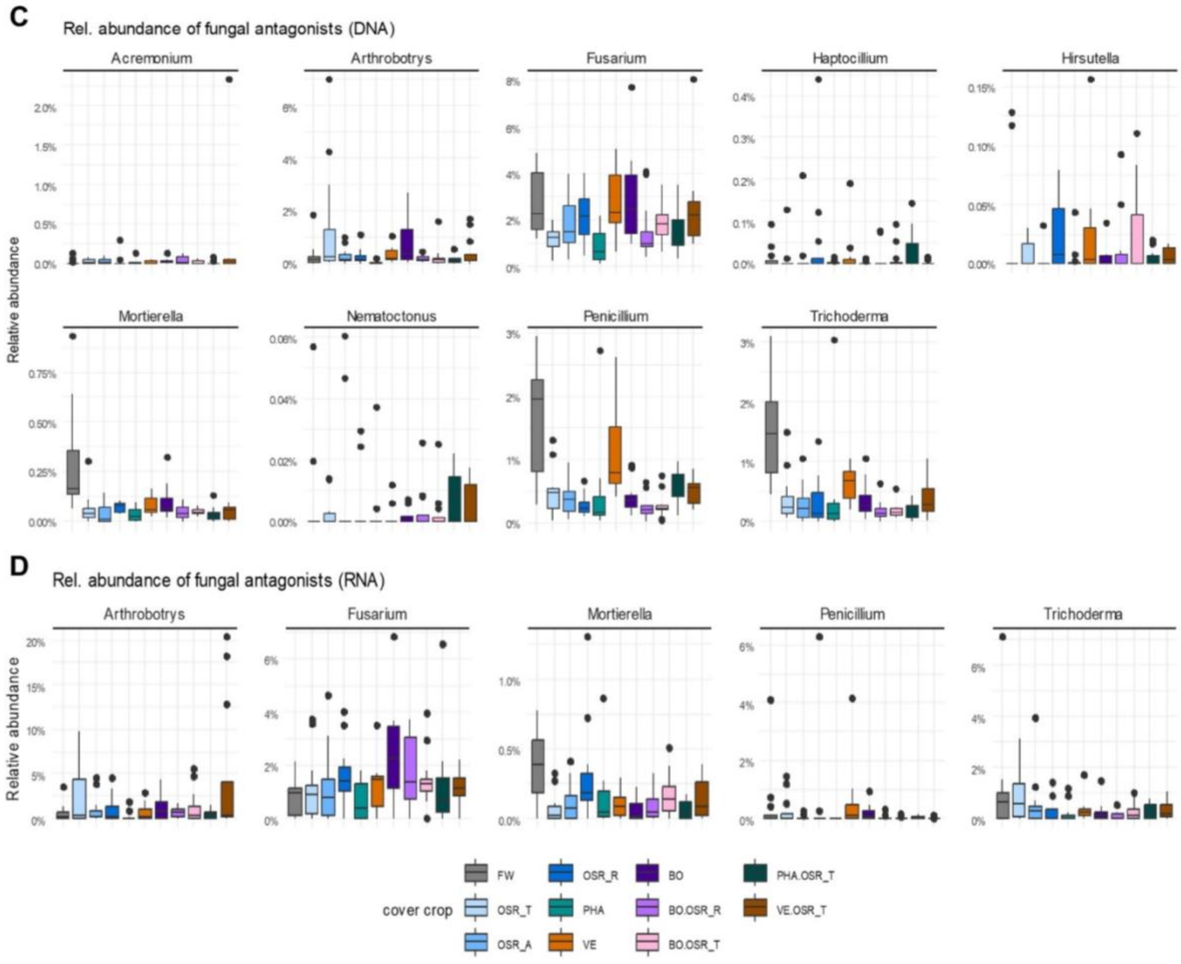
